# Restoring Glutamate receptosome dynamics at synapses rescues Autism-like deficits in Shank3-deficient mice

**DOI:** 10.1101/2020.12.30.424827

**Authors:** Enora Moutin, Sophie Sakkaki, Vincent Compan, Nathalie Bouquier, Federica Giona, Julie Areias, Elise Goyet, Anne-Laure Hemonnot-Girard, Vincent Seube, Nathan Benac, Yan Chastagnier, Fabrice Raynaud, Etienne Audinat, Laurent Groc, Tangui Maurice, Carlo Sala, Chiara Verpelli, Julie Perroy

**Affiliations:** IGF, University of Montpellier, CNRS, INSERM, Montpellier, France; Cnr Neuroscience Institute, Via Vanvitelli 32, 3220129 Milan, Italy; Interdisciplinary Institute for NeuroScience, CNRS, UMR 5297, Centre Broca Nouvelle-Aquitaine, 146, rue Léo-Saignat, 33076 Bordeaux, France; PhyMedExp, Univ Montpellier, INSERM, CNRS, CHU de Montpellier, France; MMDN, Univ Montpellier, EPHE, INSERM, Montpellier, France

**Author notes:** equal contribution (Sophie Sakkaki, Vincent Compan, Nathalie Bouquier).

## Abstract

Shank3 monogenic mutations lead to Autism Spectrum Disorders (ASD). Shank3 is part of the glutamate receptosome that physically links ionotropic NMDA receptors to metabotropic mGlu5 receptors through interactions with scaffolding proteins PSD95-GKAP-Shank3-Homer. A main physiological function of the glutamate receptosome is to control NMDA synaptic function that is required for plasticity induction. Intact glutamate receptosome supports glutamate receptors activation and plasticity induction, while glutamate receptosome disruption blocks receptors activity, preventing the induction of subsequent plasticity. Despite possible impact on metaplasticity and cognitive behaviors, scaffold interaction dynamics and their consequences are poorly defined. Here we used mGlu5-Homer interaction as a biosensor of glutamate receptosome integrity to report changes of NMDA synaptic function. Combining BRET imaging and electrophysiology, we show that a transient neuronal depolarization inducing NMDA-dependent plasticity disrupts glutamate receptosome in a long-lasting manner at synapses and induces signaling required for the expression of the initiated neuronal plasticity such as ERK and mTOR pathways. Glutamate receptosome disruption also decreases NMDA/AMPA currents ratio, freezing the sensitivity of the synapse to subsequent changes of neuronal activity. These data show the importance of a fine-tuning of protein-protein interactions within glutamate receptosome, driven by changes of neuronal activity, to control plasticity. In a mouse model of ASD, a truncated mutant form of Shank3 prevents the integrity of the glutamate receptosome. These mice display altered plasticity, anxiety-like and stereotyped behaviors. Interestingly, repairing the integrity of glutamate receptosome and its sensitivity to the neuronal activity rescued synaptic transmission, plasticity and some behavioral traits of Shank3∆C mice. Altogether, our findings characterize mechanisms by which Shank3 mutations cause ASD and highlight scaffold dynamics as new therapeutic target.

## Introduction

Autism Spectrum Disorders (ASD) constitute a group of neurodevelopmental disorders characterized by deficits in social communication and stereotyped behaviors such as restricted and repetitive behaviors, interests or activities (DSM5 https://doi.org/10.1176/appi.books.9780890425596). At least 25% of ASD have a genetic etiology and many of the associated genes involved regulate synaptic plasticity, leading to the “synaptic hypothesis”. According to this hypothesis, ASD would be the consequence of synaptic impairment(1). Mutations in the SHANK3 gene are one of the most common genetic causes of ASD, grouped together under the term of “Shankopathies”(2–5). Many mouse models of Shankopathies exist and recapitulate cognitive deficits found in patients, making them relevant to study the synaptic hypothesis. Shank3-deficient mice exhibit molecular impairments causing deficits in NMDA receptor-mediated neurotransmission associated with cognitive deficits(6–10) but little is known about how mutations in SHANK3 contribute to these dysfunctions. To tackle this issue, we used Shank3ΔC mice. Those mice have a disruption of Shank3 gene at exon 21, which revealed haplo-insufficiency of Shank3 gene as a causal mechanism of ASD in humans(11) and also triggered related phenotypes in Shank3ΔC mice(7). We here explored the molecular dynamics of Shank3 complex at synapses following neuronal excitation in WT and Shank3ΔC mice and its relevance for the control of glutamate receptors function, plasticity and behavior. In addition, we devised a strategy to rescue synaptic functioning in Shank3ΔC mice.

The postsynaptic density (PSD) of excitatory synapses is complex and dynamic in composition and regulation(12). Shank3 appears as the centerpiece of the PSD. This scaffolding protein contains an N-terminal ankyrin repeat domain, a SH3 domain, a PDZ domain, a proline-rich domain and a C-terminal SAM domain(13,14). By linking together major actors of the PSD, Shank3 is deeply involved in the organization and function of the glutamatergic post-synapse(15–17). It physically interacts with the C-terminal domain of GKAP through its PDZ domain and with Homer EVH1 (Ena/Vasp homology) N-terminal domain through its proline-rich domain. Thereby, it creates a physical molecular bridge between ionotropic NMDA and metabotropic mGlu5 receptors through the PSD95-GKAP-Shank-Homer complex, that we here call the “glutamate receptosome” (Figure 1A). These receptor-associated scaffolding proteins are much more than just placeholders since they can modulate receptors expression and activity, and therefore modulate their associated signaling pathways (see (18,19) for reviews). Synaptic protein interaction network undergoes input-dependent rearrangement by neuronal activity(20). In this context, it’s the dynamics of the scaffolding complex that play the leading role in the control of receptor functioning and synaptic transmission. Because mGlu5 and NMDA receptors play tremendous functions in the induction of synaptic plasticity, fine regulations of scaffold interactions within the glutamate receptosome finely tune the availability of synapses for plasticity induction(21). A noteworthy example of these molecular dynamics is the one controlled by Homer isoforms(22). The constitutive, long forms of Homer ensure the glutamate receptosome integrity by linking Shank3 to mGlu5 receptor(23) through the multimerization of Homer(24). By opposition, a short monomeric form of Homer, Homer1a, encoded by an immediate early gene after a sustained neuronal activity, competes with the multimeric form of Homer to interact with mGlu5 receptor, freeing it from the complex(22). When the scaffolding complex is disrupted, mGlu5 and NMDA receptors can directly interact, resulting in a reciprocal functional inhibition(21,25). This restructuration occurs after sustained neuronal activity (like those required to induce long term potentiation) and serves as negative feedback to decrease synaptic NMDA receptors activity. As a consequence, the efficacy of synaptic transmission is locked, preventing further induction of synaptic plasticity (Figure 1A). However, very little is known about the properties of these molecular dynamics at synapses to promote metaplasticity. The question of how these scaffolds remodel over time, in response to neuronal activity, remains open. A better understanding of this molecular tinkering would enable to repair deficient interactions in autism models.

**Figure 1:**
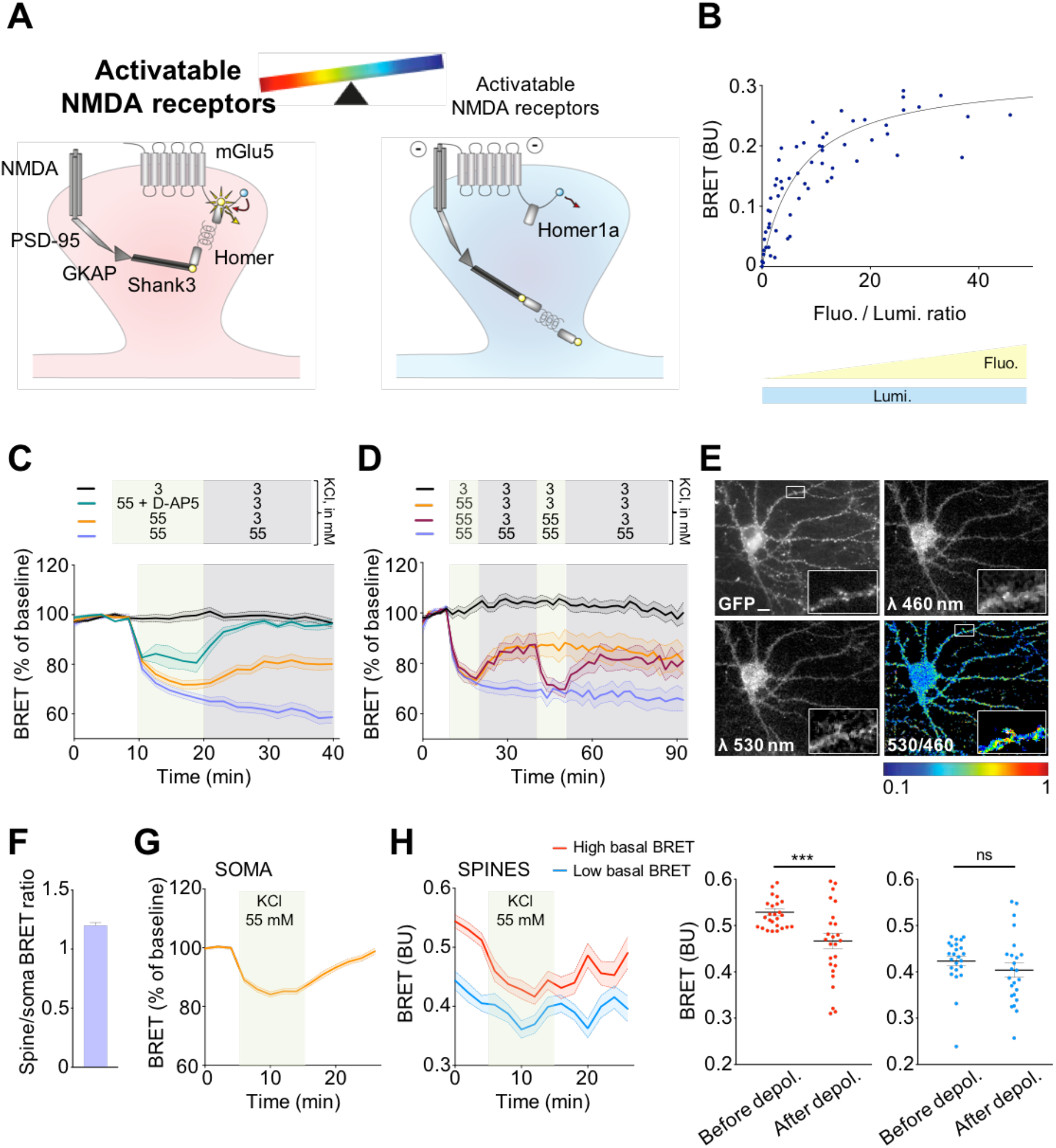
Transient depolarization induces a long-lasting disruption of mGlu5-Homer interaction in spines. **A:** Schematic illustrations of the glutamate receptosome organization in spines from WT mice. BRET intensities between mGlu5-NLuc and Venus-Homer enables to report intact (left) or disrupted (right) complexes, which are respectively highlighting synapses that are available or not for plasticity induction. **B-H:** BRET recordings between mGlu5-NLuc and Venus-Homer in neurons from hippocampal cultures in cell population (**B-D**) or in microscopy (**E-H**). BU stands for BRET units. **B:** Individual readings obtained from 5 independent experiments were pooled. Curves were fitted using a nonlinear regression equation, assuming a single binding site (GraphPadPrism version 7). **C, D, G, H:** Real time measurements of BRET variations measured in neurons during changes of membrane potential with KCl 3 and 55 mM. Data are mean ± SEM of BRET intensities recorded in triplicate from at least 3 independent experiments (**C** and **D**) or on 18 to 45 neurons in at least 6 independent experiments (**G** and **H**). In **C,** green circles, D-AP5 100 μM was added during the transient depolarization. **E:** Example of BRET imaging recorded by videomicroscopy in one neuron. The pictures show expression of Venus-Homer1c (GFP), mGlu5-NLuc (Em 460 nm), Venus-Homer1c excited by the energy transfer coming from mGlu5-NLuc (Em 530 nm) and the BRET signal (530/460 ratio). Scale bar 10 μm. **F:** Ratio of BRET intensities recorded in spines over soma on 45 neurons from 8 independent cultures. **H**: Left panel, BRET variations recorded on 50 dendritic spines segregated according to the intensity of their basal BRET: higher (red), or lower (blue) than 0.48, the median BRET value measured on the 3 first time-points. Right panels, BRET intensities in spines measured on the 3 first (before depol.) or 3 last (after depol.) time-points in spines with high (red) or low (blue) basal BRET. ns, *** indicate p-value > 0.05, < 0.001, respectively; Paired t-test. Point-by-point statistical tests and post hoc comparisons for **C, D, G and left H** are indicated in Supplemental table 1.

Here, we used mGlu5-Homer interaction as a biosensor of the glutamate receptosome integrity in WT mice to report the spatio-temporal changes of synaptic NMDA functioning, in living cells. We found that a NMDA-dependent transient increase in neuronal activity disrupts mGlu5-Homer interaction in a long-lasting manner at synapses, inducing signaling involved in plasticity like ERK and mTOR pathway activation. Scaffold disruption also triggers feedback inhibition of NMDA/AMPA currents ratio. Hence, disrupting mGlu5-Homer interaction before plasticity induction abolished ERK/mTOR activity-dependent signaling, highlighting the role of scaffold remodeling in metaplasticity. These last conditions were similar to those that apply to Shank3ΔC mice: the glutamate receptosome is atypically disrupted by the lack of interaction between the truncated form of Shank3 and Homer, furthermore NMDA/AMPA currents ratio is decreased compared to WT mice. However, by expressing a molecular bridge that restores glutamate receptosome integrity in Shank3ΔC mice and its sensitivity to neuronal activity, we could recover NMDA/AMPA ratio, rescue ERK and mTOR signaling and improve autistic-like behaviors such as anxiety and repetitive stereotyped behavior. Modulation of glutamate receptosome integrity could therefore represent a new therapeutic strategy in Shankopathies and more largely in ASD.

## Materials and methods

### Animal Handling

All animal procedures were conducted in accordance with the European Communities Council Directive, supervised by the IGF institute’s local Animal Welfare Unit (A34-172-41) and approved by the French Ministry of research (agreement numbers: APAFIS#23357-2019112715218160 v4 and APAFIS#23476-2020010613546503 v4). The Shank3ΔC mice have a deletion of exon 21 that includes the Homer-binding domain (Jackson laboratory, Bar Harbor, ME) and were backcrossed to C57BL6/J mice. Heterozygous mice were bred and all mice were grouped after weaning with respect to their sex. For experiments, we used homozygous Shank3ΔC mice and WT littermates.

### Solutions used for experiments

KCl 3 mM (in mM): NaCl 140, CaCl_2_ 2, KCl 3, Hepes 10, D-glucose 10, Glycine 0.01, Bicuculine 0.01, 0.0003 Tetrodotoxin, pH 7.4 with an osmolarity around 315-320mOsm. Membrane potential was calculated using the voltage Goldman-Hodgkin Katz (GHK) equation. This equation takes into consideration two main factors: (1) the membrane permeability and (2) concentration gradient of the most abundant permeable ions, Na^+^, K^+^, and Cl^−^.

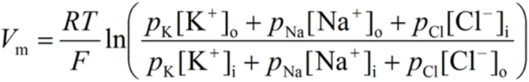

***V***_**m**_ = membrane potential.

***R*** = universal gas constant (8.314 J.K^−1^.mol^−1^)

***T*** = temperature in Kelvin (K = °C + 273.15)

***F*** = Faraday’s constant (96485 C.mol^−1^).

***p***_**K**_ = membrane permeability for K^+^. *p*_K_ : *p*_Na_ : *p*_Cl_ = 1 : 0.05 : 0.45.

**[K**^+^]_**o**_ is the concentration of K^+^ in the extracellular solution.

**[K**^+^]_**i**_ is the concentration of K^+^ in the intracellular solution.

According to GHK equation, neurons in KCl 3 mM present a membrane potential of ‒71 mV. KCl 55mM-evoked depolarization solution (in mM): NaCl 88, CaCl_2_ 2, KCl 55, Hepes 10, D-glucose 10, Glycine 0.01, Bicuculine 0.01, 0.0003 Tetrodotoxin, pH 7.4 with an osmolarity around 315-320 mOsm. According to GHK equation, this KCl 55 mM-containing solution raises membrane potential to ‒30 mV.

### Plasmids, BRET biosensors and TAT peptides

Packaging plasmid pMD2.G and psPAX2 plasmids were given by Didier Trono (Addgene plasmid #12259 and #12260).

For transgene expression, backbones of pWPT-Camk2αprom-mGlu5-NLuc and pWPT-Camk2αprom-Venus-Homer1C were all derived from pWPT-GFP plasmid (Addgene plasmid #12255). pAAV-CW3SL-mGlu5-NLuc was derived from pAAV-CW3SL-eGFP (given by Bong-Kiun Kaang, Addgene plasmid # 61463). All these plasmids were produced by Gibson Assembly (New England Biolabs) after amplification by PCR of mGlu5, Venus, Homer1C or CamK2α promoter. pAAV-CW3SL-Venus-Homer1b-GluN2B-Ctail was obtained by subcloning the Venus sequence from pAAV-CW3SL-Venus-Homer1C and the Homer1b-GluN2BCtail sequence from pGW1-GFP-Homer1b-GluN2B-Ctail into pAAV-CW3SL-mGlu5-NLuc using Gibson Assembly protocol. pGW1-GFP-Homer1b-NR2B-Ctail was built by adding the last eight AA of GluN2B (LSSIESDV) spaced by 10 Glycine to the C-terminal of Homer1b protein.

YEN BRET biosensor, that reports ERK activity, was already described(26) and subcloned in a pAAV backbone to obtain pAAV-hSyn-YEN. AIMTOR BRET biosensor, that reports mTORC1 activity, was already described(27) (pAAV-hSyn-T757-AIMTOR).

Cell-permeable TAT-fused peptides TAT C-tail (YGRKKRRQRRR-ALTPPSPFR) and the control TAT (TAT ctrl : YGRKKRRQRRR-ALTPLSPRR)(28) were synthesized by Smart Bioscience company, resuspended in H_2_O at a concentration of 1 mM and stored at −80 °C. Please note that TAT C-tail can disrupt mGlu5-Homer but also Shank3-Homer interaction as it mimics the proline-rich EVH1 binding domain that is present in mGlu5 and Shank3. TATs were used at 5 μM for BRET experiments and at 10 μM for electrophysiology.

### HEK293T culture

HEK293T cells were cultured in DMEM (Invitrogen) containing 10% FBS and antibiotics (Penicillin/Streptomycin).

### Viral productions

Lentiviral particles were produced as described(29). Briefly, HEK293T cells were co-transfected with the targeted vector plasmid, the packaging plasmid pMD2G and the envelope plasmid pSPAX2 using phosphate calcium technique. After 3 days of incubation, the cell supernatant was filtered and incubated overnight with polyethylene glycol to allow lentivirus concentration by precipitation. After centrifugation, pellets containing lentiviral particles were resuspended in PBS, aliquoted and stored at −80 °C.

AAV-DJ particles were produced using a commercial kit, according to the manufacturer’s protocol (AAV purification kit, reference 6666 from Takara and reference VPK-140 from Cell Biolabs). AAV particles were resuspended in PBS with an approximate titer of 1 x 10^14^ genome copies/ml, aliquoted and stored at −80 °C.

### Hippocampal primary cell culture and transduction

For the majority of the experiments, hippocampal neuronal primary cultures were prepared from P0-P2 mice as described(29). Briefly, brain hippocampi were mechanically and enzymatically dissociated with papain (Sigma-aldrich) and hippocampal cells were seeded in Neurobasal-A medium (Gibco) supplemented with B-27 (Gibco), Glutamax (Gibco), L-glutamine (Gibco), antibiotics (Gibco) and Fetal Bovine Serum (Gibco). After 2 days in culture, Cytosine β-D-arabinofuranoside hydrochloride (Sigma-aldrich) was added to curb glia proliferation. The day after, 75% of the medium was replaced by BrainPhys medium (Stemcell technologies) supplemented with B-27 (Gibco), Glutamax (Gibco), and antibiotics (Gibco). At DIV6, 15% of the medium was replaced by fresh medium containing lentiviral particles or AAV particles for neuronal transduction, as recently described(29). For immunocytochemistry and for single particle tracking experiments hippocampal neuronal primary cultures were prepared from embryonic rats (E18). Briefly, brain hippocampi were mechanically and enzymatically dissociated with trypsin (Sigma-aldrich) and hippocampal cells were seeded in Neurobasal medium (Gibco) supplemented as for post-natal cultures. After 3 days in culture, Cytosine β-D-arabinofuranoside hydrochloride (Sigma-aldrich) was added only for cultures from mice.

### Co-immunoprecipitation

Hippocampi from 14 Shank3ΔC mice and 14 WT littermates 2 to 4-months old were solubilized in cold lysis buffer (in mM): Tris-HCl (50), NaCl (120), NaF (50), Triton 1% supplemented with protease and phosphatase inhibitors (Thermofisher). After 10 up and down using Dounce homogenizer, samples were kept on ice for 30 min and centrifuged at 15.900g for 10 min at 4°C. Concentration of solubilized proteins was quantified using BCA assay (Sigma-aldrich). All samples were then set at the same volume and concentration for the rest of the protocol. One equal volume was kept for total extracts and the rest was incubated with antibody against mGlu5 (rabbit clone D6E7B from Cell Signaling) with rotation overnight at 4 °C. Then, protein A/G magnetic beads (Pierce, Life technologies) were added in samples for 4 hours with rotation at 4 °C. After washings, proteins were eluted in LDS Sample buffer (Thermofisher) supplemented with 2% ß-mercaptoethanol, resolved on 4-12% gradient NuPAGE Bis-Tris gels (Thermofisher), transferred onto nitrocellulose membranes and detected by immunoblot using the following primary antibodies: mGlu5 (rabbit clone D6E7B from Cell Signaling, dilution 1:1000), actin (mouse clone from DSHB, reference JLA20, dilution 1:2000), GluN1 (rabbit clone D65B7 from Cell Signaling, dilution 1:1000).

### BRET measurements

Just before BRET recordings, the culture medium was replaced by a KCl 3 mM solution. To assess the effects of depolarization on BRET signal, KCl 3 mM was replaced by a KCl 55mM solution. KCl3 and 55 mM solutions are described above.

**Cell population BRET measurements** were performed using the Infinite F500 (TECAN) 96-well plate-reader on DIV 14 to 16 neurons previously transduced at DIV6 with pWPT-Camk2αprom-mGlu5-NLuc and pWPT-Camk2αprom-Venus-Homer1C or with pAAV-CW3SL-mGlu5-NLuc and pAAV-CW3SL-Venus-Homer1b-GluN2B-Ctail. BRET computed as the ratio of the intensity detected from the Venus acceptor entity (520-575nm band-pass filter, Em530) to the NLuc donor (390-450nm band-pass filter, Em460), after the addition of 20 μM Furimazine. BRET saturation was measured in neurons transduced with constant concentrations of mGlu5-NLuc and increasing concentrations of Venus-Homer1c. BRET was measured in each well every 2 minutes (3 wells per condition in each experiment, pooled for the analysis).

**Single-cell BRET imaging** was performed as previously described(26,30) on hippocampal neurons from Shank3ΔC or WT mice. Neurons cultured on glass-bottom culture dishes (reference 81156-400, Ibidi) were transduced at DIV 6 with pWPT-Camk2αprom-mGlu5-NLuc and pWPT-Camk2αprom-Venus-Homer1C or with pAAV-hSyn-T757-AIMTOR or with pAAV-hSyn-YEN and/or with pCW3SL-Venus-Homer1b-GluN2B-Ctail. We performed acquisitions between DIV13 and DIV15 for experiments on ERK and mTOR and between DIV15 and DIV17 for experiments on mGlu5-Homer1 interaction. All images were obtained using a bioluminescence-dedicated inverted fluorescence microscope (Axiovert 200M, Carl Zeiss) with a Plan Apochromat 63× /1.40 oil M27 objective at room temperature and collected with an evolve camera (Photometrics) equipped with an EMCCD detector, back-illuminated, On-chip Multiplication Gain. Sequential acquisitions of light emission at 460 nm and 530 nm were performed at 5 MHz, Em gain 200 and conversion gain 3e^−^/ADU, binning 1, with emission filters FF01-450/70-25 (Semrock) and HQ535/50 nm (No. 63944, Chroma), respectively. Furimazine (50 μM) was applied 1 minute before acquisition and the 530 nm/460 nm ratio was read every two minutes for each neuron. For BRET between mGlu5 and Homer and for AIMTOR biosensor, we sequentially exposed the donor 10 s and the acceptor 20 s. For ERK biosensor, we sequentially exposed the donor 5 s and the acceptor 10 s. BRET was measured in each field every 2 or 3 minutes. Analysis was performed as previously described(31). Briefly, we used an open source toolset for Fiji (https://github.com/ychastagnier/BRET-Analyzer) that performs 4 key steps: (1) image background subtraction, (2) image alignment over time, (3) a composite thresholding method of the image and (4) pixel by pixel division of the image and distribution of the ratio intensity on a pseudocolor scale.

### AAV injections and immunohistochemistry

We injected Shank3ΔC and WT male mice littermates between 6 and 8 post-natal days. Mice were anesthetized with isoflurane (Vetflurane®, 2-5%) and AAV vectors were injected into the lateral ventricles randomly across the different litters. 0.5 μl were injected through each injection site at the following coordinates respective to bregma: Antero-Posterior: −2.5 mm, Median-Lateral: −2.5 mm/+2.5 mm, Dorso-Ventral: −1.25 mm based on Allen Developing Mouse Brain Atlas (2008). After weaning, mice were divided in 3 groups and kept in separate cages, WT injected with AAV containing eGFP, Shank3ΔC injected with AAV containing eGFP and Shank3ΔC injected with AAV containing Homer-GluN2B chimaera. Generalized Estimated Equation (GEE), with group and cage as predictors and each specific behavioral measurement as variable, ruled out potential cage effect. To evaluate the efficiency of the viral infection, mice brains were kept after behavioral experiments in 4% paraformaldehyde during 24 h, washed and then sliced into coronal sections of 30 μm thickness using a vibratome. Sections were processed for immunofluorescence to enhance eGFP signal. Briefly, slices were permeabilized in PBS containing triton 0.2%, then blocked with PBS containing BSA 3% and incubated overnight with an antibody against eGFP (chicken, AVES, GFP-1020) diluted in PBS containing BSA 1% and triton 0.1% and then incubated with appropriate secondary antibody and Hoechst for nuclei staining. Images were acquired using an AxioImager Z1 Zeiss microscope equipped for optical sectioning (Apotome) and with appropriate epifluorescence filters. Supplemental Figure 7 shows transgene expression in brain areas.

### Electrophysiology in acute slices

Shank3ΔC mice or WT littermates aged between postnatal day 14 and 20 were briefly anesthetized with isoflurane and decapitated. Brains were quickly removed in oxygenated (5% CO_2_ and 95% CO_2_) ice-cold extracellular solution containing (in mM): Sucrose (215), D-glucose (20), Sodium pyruvate (5), KCl (2.5), NaH_2_PO_4_ (1.25), NaHCO_3_ (25.9), MgCl_2_ (7), CaCl_2_ (1) (pH 7.4, 315mOsm). Acute coronal slices 350 μm thick were cut using a Leica VT1200S vibratome in the same extracellular solution and then incubated at 33°C in ACSF (in mM): NaCl (126), D-glucose (20), Sodium pyruvate (5), KCl (2.5), NaH_2_PO_4_ (1.25), NaHCO_3_ (25.9), MgCl_2_ (1), CaCl_2_ (2) (pH 7.4, 310 mOsm). Extracellular solution temperature was then slowly cooled until reaching room temperature and slices were maintained in this oxygenated solution until recordings (0.5-5 h). Before recordings, a cut was made between CA3 and CA1 to prevent the propagation to CA1 of CA3-generated bursts of action potentials upon blockade of GABA-A receptors. For recordings, slices were transferred in a recording chamber, perfused with ACSF (2-3 ml/min) at room temperature. CA1 pyramidal cells were visually selected using infrared LED illumination (Scientifica Ltd) and an Olympus 40x water immersion objective. For experiments on mice injected with viruses, fluorescent CA1 pyramidal neurons were selected using a 480nm LED (Cairn Research) on the epifluorescent port of the microscope. A patch pipette filled with ACSF (1 MΩ) placed in the stratum radiatum was used as a monopolar stimulating electrode. The recording electrode (2-3 MΩ) was filled with the following intracellular medium (in mM): CsMeSO_3_ (125), Hepes (10), EGTA (10), TEA-Cl (8), 4-aminopyridine (5), GTP-Na (0.4), ATP-Na_2_ (4), CaCl_2_ (1) MgCl_2_ (1) (pH 7.4, 288 mOsm). Whole-cell patch-clamp recordings were obtained using Axopatch 700B (Molecular Devices) and performed in the presence of 10 μM bicuculine to block fast inhibitory transmission. Recordings were filtered at 1 kHz and digitized at 10 kHz. Paired EPSCs (100 ms interval) were evoked at 0.1 Hz in neurons held in voltage clamp at – 80 mV to measure AMPAR-mediated EPSCs and then at + 40 mV for NMDAR-mediated EPSCs.

Raw data were analyzed using Clampfit (pClamp software version 10.7, Molecular Devices). The amplitude of EPSCs was quantified from the average of 15 to 30 consecutive responses. The paired-pulse ratio was calculated as the average peak amplitude of the second response divided by that of the first one.

### Behavioral experiments

Behavioral experiments were performed on the injected animals between 7-20 weeks old (see AAV injections and immunohistochemistry section for group details). Mice were housed in group-cages with 3-6 individuals per cage. Cages were supplemented with minimal enrichment (cotton nestlets). Mice were accustomed to the experimental animal facility and were manipulated every day by the experimenter for at least one week before behavioral testing. Behavioral experiments were performed between 8:00 am–4:00 pm in a light color wall room with light devices. Level of illumination used was as described in Kouser et al(7). The experimenter was blind to the genotype and to the material injected during testing.

**Open Field** consisted of a white square arena (50 cm x 50 cm). Mice were placed in the center of the arena and left to explore freely for 10 min. Total distance travelled, speed and time spend in the center zone (defined as a 35 cm side square) were measured during 10 minutes by video tracking (Viewpoint, Lissieu, France).

#### Self-grooming

Mice were placed in a 25 cm Plexiglas cylinder and video monitored. Time spend self-grooming was visually measured during 10 min.

#### Marble burying

Mice were placed individually in a clean cage with 10 cm thick bedding together with 15 marbles evenly distributed on top of the bedding. After 30 min in the cage, mice were removed and we evaluated the level of marble burying according to the following Score: 0 for a totally buried marble; 1 for a half-buried marble and 2 for a totally visible marble.

#### Rotarod

Mice were put on an accelerating rotarod apparatus, 2 to 45 rpm in 5 min (Ugo Basile, Gemonio, Italy). Time to fall was measured during 4 sessions spaced 30 minutes apart and the average of the 3 last trials was measured.

#### Auditory Fear conditioning

We performed an auditory fear conditioning protocol during two consecutive days. The experiments were carried out in a fear conditioning apparatus comprising a test box (20 cm width × 20 cm length × 20 cm height) placed within a sound proof chamber (Panlab, Harvard Apparatus). Two different contextual configurations were used (A: square configuration, white walls, metal grid floor, washed with 70% ethanol; B: square configuration, black walls, white floor, washed with 1% acetic acid). On day 1, mice were subjected to a habituation session in context A. After 10 min of habituation to the box, mice received 5 pairing of one tone (CS+: 2.5 kHz, 80 dB, 10 s) with an unconditioning stimulus (US: 0.6 mA scrambled footshock, 2 s, coinciding with the last 2 s of CS+ presentation). On day 2, test day, mice were subjected to a 10 min habituation in context B and then received 12 presentations of the CS+ alone. Freezing behavior during CS+ presentations was analyzed using a load cell coupler (Panlab, Barcelona, Spain) and was defined as the lack of activity above a calibrated threshold for a duration of 2 s or more as determined with the Freezing software (Panlab). The average time spent freezing prior to presentation of the sounds (habituation in context B) during the test sessions was used as a measure for contextual fear generalization. We computed a ratio as the fraction of time spent freezing during the 30 seconds following CS+ presentation over the fraction of contextual time freezing.

### Statistical Analysis

Statistical analysis was carried out using GraphPad Prism 7 software (GraphPad Software Inc.). Statistical tests and post hoc comparisons are indicated in the figure legends and in supplemental Table 1.

## Results

### Transient increase of neuronal activity induces a long-lasting disruption of glutamate receptosome in spines

mGlu5-Homer interaction reports synaptic availability for plasticity induction. The loss of mGlu5-Homer interaction, by preventing mGlu5 and NMDA receptors activation(21, 25), highlights synapses in which induction of plasticity is abolished (Figure 1A). Due to recent advances in BRET technology(26, 31), we followed mGlu5-Homer interaction in real time in living hippocampal neurons, to report NMDA synaptic function and tag synapses that are available for plasticity induction. We fused mGlu5 to the energy donor Nano-Luciferase (NLuc) and Homer with the fluorescent acceptor Venus. Under stable mGlu5-NLuc expression, the BRET signal increased hyperbolically as a function of the Venus-Homer expression level, indicating a specific interaction between mGlu5 and Homer (Figure 1B). This BRET signal remained stable over time (Figure 1C, black, control condition). Neuronal activity modulates the establishment and refinement of neuronal connections, mainly through its effects on dendritic morphology and synaptic plasticity. To determine the effect of neuronal activity on mGlu5-Homer interaction, we treated primary hippocampal cultures with 55 mM KCl to allow a systematic, well-controlled and long-lasting depolarization of neurons. Neuronal stimulation with KCl causes an estimated depolarization of – 30 mV (see Materials and methods), activation of voltage-gated receptors, and induction of immediate-early genes (IEGs) simulating physiological stimulation(32). We confirmed IEGs expression induced by this protocol, as illustrated by Homer1a expression (Suppl. Figure 1). In neuronal population, depolarization caused a fast decrease of the BRET signal intensity between mGlu5-NLuc and Venus-Homer (Figure 1C, purple). Alternative protocols to induce plasticity, like direct stimulation of NMDA receptors, also disrupted mGlu5-Homer interaction (Suppl. Figure 2). Setting back neurons to their resting membrane potential lead to a partial recovery of the BRET signal (Figure 1C, orange). Thus, mGlu5-Homer interaction is activity-dependent and only partially reversible upon repolarization. A second subsequent transient increase in neuronal activity fully and reversibly disrupted the recovered mGlu5-Homer complexes (Figure 1D, red). These data defined two patterns of mGlu5-Homer interaction: one stably disrupted by transient depolarization; a second one highly reversible, which disruption coincides with the cell membrane depolarization. To undergo long-lasting disruption, the first pool of mGlu5-Homer interaction required NMDA receptor activation. In presence of NMDA receptor antagonist, D-AP5, mGlu5-Homer interaction was indeed disrupted by transient depolarization, in a fully reversible manner (Figure 1C, green). This dichotomy of NMDA receptor involvement, plus the fact that the expression of the glutamate receptosome components, are confined to the PSD, favored the hypothesis of regional segregation of the mGlu5-Homer interaction dynamics. In support of this hypothesis, BRET imaging demonstrated sub-cellular specific patterns for mGlu5-Homer interaction dynamics in spines compared to soma (Figure 1E-H). Images in figure 1E show mGlu5-NLuc expression (460 nm), Venus-Homer expression (GFP) and the intensities of BRET signal in subcellular compartments (530/460). BRET signal was indeed 1.2-fold stronger in spines than in soma (Figure 1F). In the soma, we measured a homogenous mGlu5-Homer interaction, which was disrupted by transient depolarization and fully reversed within ten minutes (Figure 1G). By opposition, we noticed an important heterogeneity between spines and divided them in two equivalent populations depending on their BRET signal intensities (Figure 1H). Importantly, the highest the BRET signal was, the strongest was the efficiency of neuronal activity to disrupt the interaction. In these spines, the transient depolarization irreversibly disrupted mGlu5-Homer interaction (Figure 1H, red). Low basal BRET signal in spines may reflect recent synaptic activation. Consistently, no significant change of BRET was recorded in these spines following transient depolarization (Figure 1H, blue). Altogether, these experiments show that the synaptic availability for plasticity induction, reported by mGlu5-Homer interaction, is heterogenous among spines and is blocked at least for 1 hour following transient depolarization.

### Glutamate receptosome disruption decreases NMDA/AMPA excitatory postsynaptic currents ratio and impairs plasticity induction

Spines containing undisrupted glutamate receptosome (high BRET, Figure 1H) undergo activity-driven NMDA receptor activation, followed by long-lasting complex disruption. Highly permeable to calcium ions, NMDA receptors play a key role in activity-induced long-term changes of synaptic strength. mGlu5 direct interaction with NMDA receptors enables mutual functional inhibition(21,25). We also know that Homer isoforms control mGlu receptors’ localization(33). Therefore, scaffolding complex disruption could enhance mGlu5 receptor mobility enabling its direct interaction with and inhibition of NMDA receptor. This was indeed supported by our previous report using single-molecule tracking, where mGlu5 was significantly more mobile at synapses in hippocampal neurons when mGlu5-Homer interaction was disrupted, causing an increased synaptic surface co-clustering of mGlu5 and NMDA receptors(34). Consistently, we found that a transient increase in neuronal activity, inducing disruption of glutamate receptosome (Figure 1), enhanced endogenous mGlu5 mobility (Suppl. Figure 3). To further understand the role of glutamate receptosome dynamics on glutamatergic synapse physiology, we disrupted mGlu5-Homer interaction and studied the functional consequences on NMDA/AMPA excitatory postsynaptic currents ratio and on the activation of ERK and mTOR pathways as readouts of synaptic plasticity. We used a cell-permeant TAT peptide containing the proline-rich motif of the mGlu5 C-terminal tail (TAT C-tail) to compete with mGlu5 to bind the EVH1 domain of Homer(28). TAT C-tail disrupted mGlu5-Homer interaction in less than 10 minutes, while a control TAT peptide containing a mutated Homer binding motif had no significant effect (Suppl. Figure 4).

The NMDA/AMPA ratio is an important parameter that strongly influences the integrative properties of excitatory synapses. Acute treatment of brain slices with the TAT C-tail (but not TAT control) induced a strong decrease of the NMDA/AMPA postsynaptic current ratio in the hippocampus CA1 (Figure 2A). Therefore, depolarization-induced glutamate receptosome disruption would increase mGlu5 mobility, leading to feedback inhibition of NMDA activity. We then studied the activation of ERK and mTORC1 pathways, in real-time, using specific intramolecular BRET-based biosensors in hippocampal neuronal cultures (YEN(26) or AIMTOR(27), respectively, Figure 2B). Transient depolarization enhanced ERK (Figure 2C, TAT ctrl) and mTOR (Figure 2D, TAT ctrl) signaling pathways. Interestingly, disrupting glutamate receptosome prevented both pathways activation by a subsequent depolarization (Figure 2C and 2D, TAT C-tail). Altogether, these results show that impaired glutamate receptosome dynamics alter mGlu5 mobility, NMDA/AMPA ratio and synaptic plasticity. These data explain how disruption of glutamate receptosome prior to changes of neuronal activity prevents NMDA-dependent plasticity induction(21). Hence, a first neuronal activation inducing NMDA-dependent plasticity, by disrupting the complex in a long-lasting manner and inducing feed-back inhibition of NMDA receptors, would freeze synaptic sensitivity to subsequent neuronal activation, allowing the expression of the initiated plasticity.

**Figure 2:**
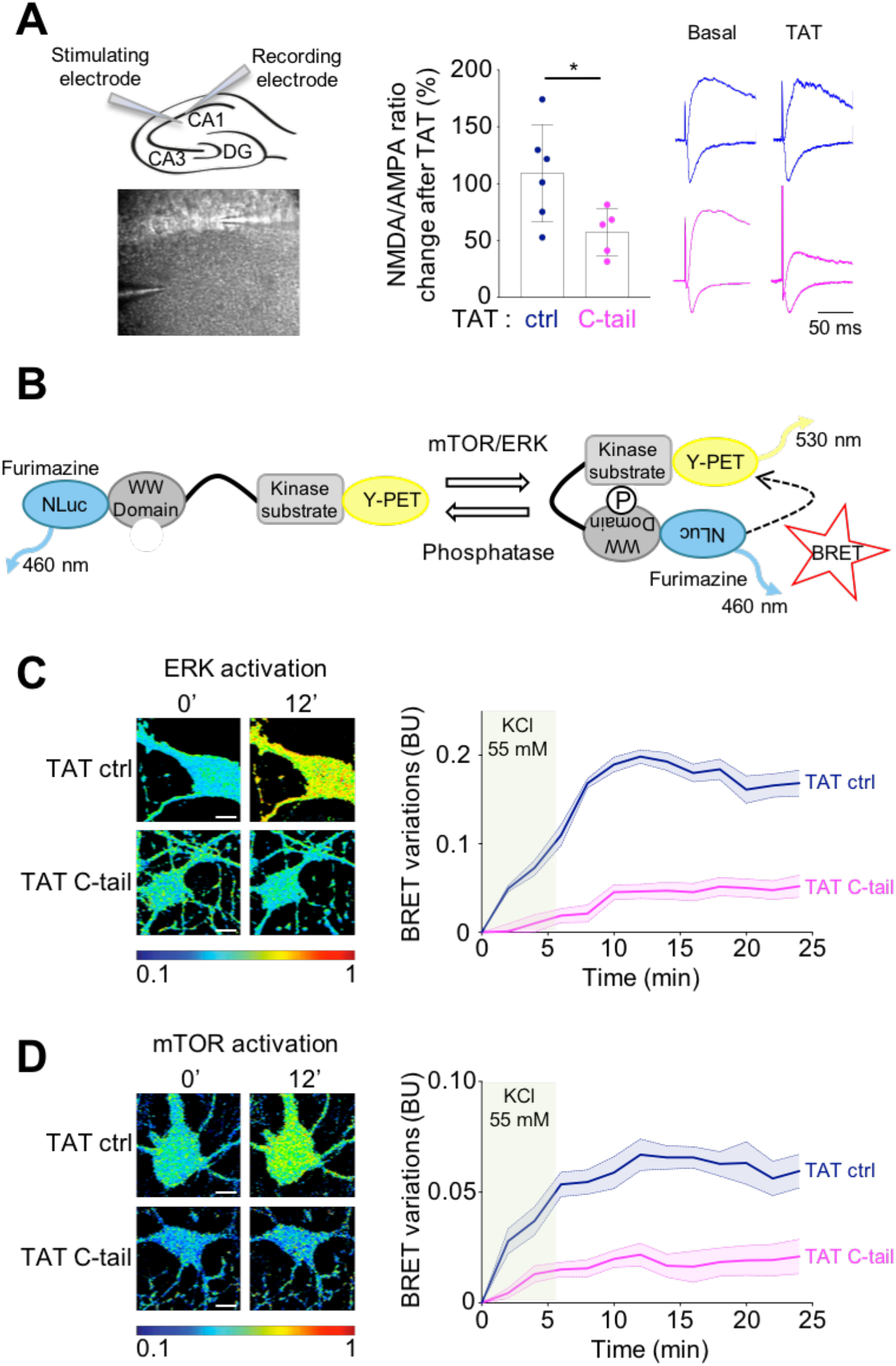
Acute mGlu5-Homer disruption impairs NMDA/AMPA postsynaptic currents ratio and ERK and mTOR signaling pathways. **A:** CA1 pyramidal neurons were clamped at −80 and +40 mV to record AMPA and NMDA currents, respectively, before and 10 min after perfusion of TATs (ctrl or C-tail). For each neuron, the NMDA/AMPA ratio is expressed as the % of NMDA/AMPA ratio before TAT perfusion. n = 5-6 neurons from 3-4 mice. Right panel: representatives average traces at −80 and +40 mV of the same recording before and 10 min after TAT perfusion. **B:** Schematic representation of the conformational changes of mTOR or ERK BRET biosensors upon their activation. **C, D:** BRET variations in neurons expressing YEN (**C**) or AIMTOR (**D**) biosensors before (0’) and 12 minutes after depolarization with KCl 55 mM (12’). Cultures were supplemented with TATs (ctrl or C-tail) 30’ before recordings (8-30 neurons, 3 independent experiments). Scale bar 10 μm. **A-D:** Data are mean ± SEM; * indicate p-value < 0.05; Mann-Whitney test. **C-D:** Point-by-point statistical tests and post hoc comparisons are indicated in Supplemental table 1.

### NMDA/AMPA excitatory postsynaptic currents ratio and neuronal activity-induced signaling are altered in Shank3ΔC mice

In Shank3ΔC mice, a truncated mutant form of Shank3 prevents the integrity of the glutamate receptosome. As glutamate receptosome remodeling after depolarization controls NMDA/AMPA postsynaptic excitatory currents ratio and ERK/mTOR activation pathways, we studied these functional aspects in Shank3ΔC mice. We observed a significant decrease of the NMDA/AMPA ratio in the CA1 area of the hippocampus of Shank3ΔC mice compared to WT mice (Figure 3A), confirming previous studies in hippocampus(7) and cortex(8). We noticed an important variability of the NMDA/AMPA ratio in WT mice, probably due to the developmental window of the experiment (P15 to P20) during which synapses are maturing while this ratio was much less variable in Shank3ΔC mice. We found no difference in the Paired pulse ratio between Shank3ΔC mice and WT mice, arguing for a postsynaptic deficit rather than presynaptic (Figure 3A). Furthermore, Shank3ΔC mice displayed strong impairment in ERK and mTOR activation by transient depolarization (Figure 3B and 3C, respectively). We have previously shown that glutamate receptosome disruption, by enabling mGlu5 and NMDA receptors interaction and mutual blockade, prevents induction of plasticity(21). Interestingly, co-immunoprecipitation experiments (Figure 3D) revealed an increased association between endogenous mGlu5 and NMDA receptors in Shank3ΔC mice hippocampi compared to WT mice. Importantly, total amount of GluN1 protein was unchanged between WT and Shank3ΔC mice (Suppl. Figure 5), as previously published(7). These results suggest that decreased NMDA/AMPA currents ratio and ERK / mTOR signaling impairments in Shank3ΔC mice are likely due to molecular dynamics deficiencies of the glutamate receptosome. Hence, we assessed mGlu5-Homer dynamics by BRET in neurons from Shank3ΔC mice compared to WT (Figure 3E). In neuronal population, we found no difference in the mGlu5-Homer disruption induced by depolarization between the two genotypes (purple curves) but a significant increase in BRET signal recovery after repolarization (orange curves) in Shank3ΔC mice compared to WT mice. Indeed, only 36.1% ± 6.9% of mGlu5-Homer disruption was long-lasting in Shank3ΔC mice versus 53.4% ± 3.9% in WT mice (Figure 3E, right panel). These results show that in Shank3ΔC mice, mGlu5-Homer interaction is still sensitive to neuronal activity but remains highly reversible, fluctuating with membrane depolarization. A second subsequent transient increase in neuronal activity indeed disrupted again mGlu5-Homer interaction in Shank3ΔC neurons (Figure 3F). These longer lasting experiments further underline the better recovery of the BRET signal in Shank3ΔC compared to WT (see Figure 1D). In addition, BRET imaging showed that more than 90% of the Shank3ΔC spines express high basal BRET values between mGlu5 and Homer (Figure 3G). In these spines, mGlu5-Homer interaction can be broken by depolarization and restored by repolarization (Figure 3H), contrary to WT spines (Figure 1H). Altogether, these experiments show that a dysregulation of mGlu5 receptosome dynamics in Shank3ΔC mice is associated with alterations of NMDA/AMPA currents ratio and ERK / mTOR signaling.

**Figure 3:**
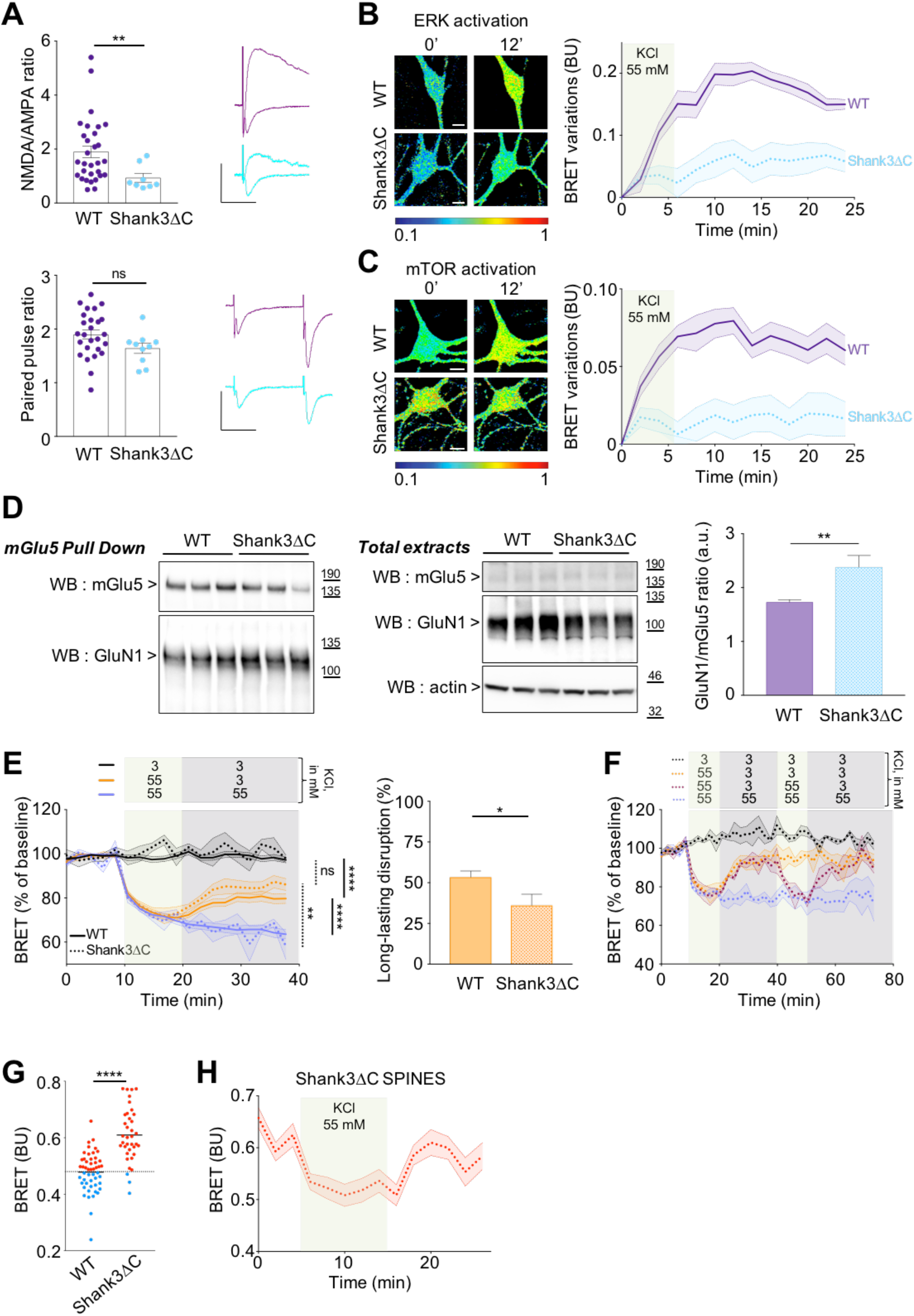
NMDA/AMPA currents ratio, ERK and mTOR signaling pathways and mGlu5-Homer interaction are altered in Shank3ΔC mice. **A:** CA1 pyramidal neurons from WT or Shank3ΔC mice were clamped at −80 and +40 mV to record AMPA and NMDA currents, respectively (100 ms interstimulus). N = 8-30 neurons from 3-15 mice. Right panels: representatives average traces at −80 and +40 mV. Scale bars 50 pA over 50 ms. **B, C:** BRET variations in WT or Shank3ΔC neurons expressing YEN (**B**) or AIMTOR (**C**) biosensors before (0’) and 12’ after depolarization (8-30 neurons, 3 independent experiments). Scale bar 10 μm. **D:** Left panel shows immunoblotting for mGlu5 and GluN1 after mGlu5 pull-down. Middle panel shows total lysates immunoblotted for mGlu5, GluN1 and actin. Graph presents the GluN1/mGlu5 ratio. **E, F, H:** BRET variations (WT = line; Shank3ΔC = dotted line) during membrane potential changes (at least 3 independent experiments). (**Left E**) One-way ANOVA multiple comparison with Tukey correction on the averaged last 2 points of each curve. Dotted/full lines to compare Shank3ΔC/WT conditions, respectively. (**Right E**) Long-lasting disruption calculated on the averaged last 2 orange time points, expressed as a % of respective ngleShank3ΔC mice. The dashed line indicates the median of basal BRET intensities in spines from WT mice (0.48). **H:** Over 37 spines (15 neurons, 3 independent experiments), the 34 having high basal BRET signal (> 0.48) are represented. Only 3 spines displayed a low basal BRET (< 0.48) and were not represented. **A-H:** Data are mean ± SEM; ns, *, **, **** indicate p-value > 0.05, < 0.05, < 0.01, < 0.0001, respectively; Mann-Whitney test (excepted for left E). Point-by-point statistical tests and post hoc comparisons for **B, C, E, F and H** are indicated in Supplemental table 1.

### Repairing scaffolding complex dynamics rescues activity-induced signaling in Shank3ΔC hippocampal neurons

In Shank3ΔC mice, the mGlu5-Homer interaction is still regulated by neuronal activity but only in a short-lived manner, having no more functional consequences on neurotransmission and plasticity. Therefore, glutamate receptosome integrity would be required for synaptic plasticity induction and expression. To test this hypothesis, we generated a molecular bridge between mGlu5 and NMDA receptors, bypassing Shank3, by fusing Homer to the C-tail PDZ ligand of GluN2B NMDA subunit after a glycine linker(35), (Figure 4A). Homer-GluN2B chimaera was correctly expressed and targeted at the spines of Shank3ΔC neurons (Figure 4B) and co-localized with PSD-95 (Suppl. Figure 6). BRET experiments in Shank3ΔC neuronal population showed that Homer-GluN2B chimaera interacts with mGlu5 in a neuronal activity-dependent mode (Figure 4C), recapitulating functional features of Homer-mGlu5 interaction in WT mice. Indeed, restoring the glutamate receptosome integrity by expressing the Homer-GluN2B chimaera rescued the long-lasting disruption induced by the neuronal activity in Shank3ΔC at 49.3% ± 9.3% (compared to 53.4% ± 3.9% in WT and 36.1% ± 6.9% in Shank3ΔC, Figure 3E). BRET experiments in Shank3ΔC neurons expressing Homer-GluN2B chimaera revealed a strong increase of ERK and mTOR activation induced by transient depolarization (Figure 4D and 4E, respectively). We noticed a significant decrease of the basal AIMTOR BRET signal intensity in Shank3ΔC neurons expressing Homer-GluN2B chimaera compared to Shank3ΔC. Thus, the inability of Shank3ΔC neurons, before rescue, to activate mTOR pathway in response to the neuronal activity could be due to mTOR hyper-activation in basal conditions (Figure 4E). Altogether, these experiments show that restoring glutamate receptosome dynamics rescued neuronal signaling and synaptic plasticity in Shank3ΔC mice. Of note, we found that in Shank3-depleted neurons using a specific shRNA, transfected Homer-GluN2B chimaera is correctly recruited to synapses while transfected WT Homer is not (Suppl. Figure 7). Hence, our strategy of bypassing Shank3 mutations/absence by reconstruction of a molecular bridge between metabotropic and ionotropic glutamate receptors with Homer-GluN2B chimaera could be extended to other Shankopathies.

**Figure 4:**
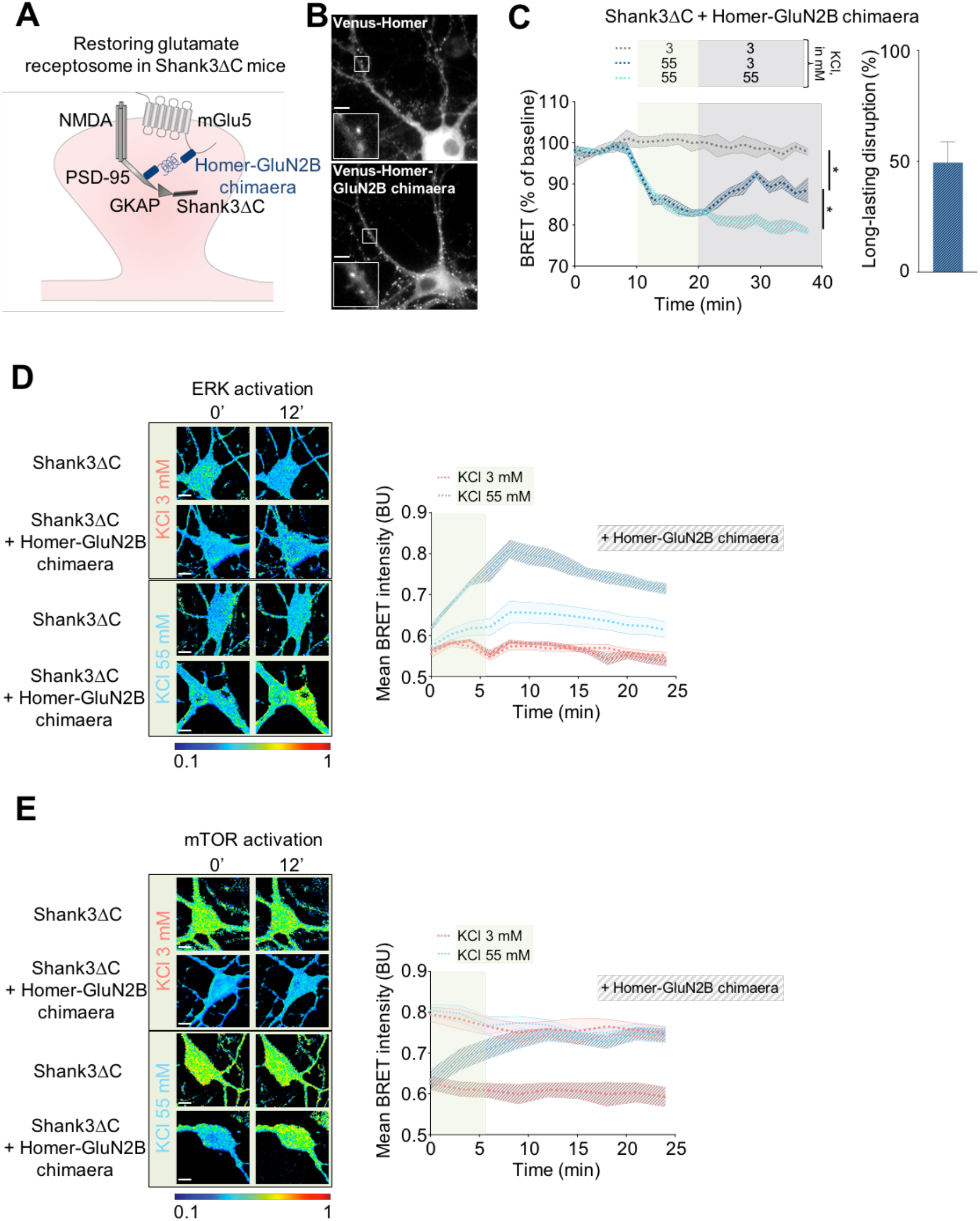
Repairing Glutamate receptosome dynamics rescues ERK and mTOR signaling pathways in Shank3ΔC mice. **A:** Schematic illustration of the glutamate receptosome reparation in Shank3ΔC neurons by expressing Homer-GluN2B chimaera to rescue synaptic availability for plasticity induction. **B:** Comparison of Homer and the Homer-GluN2B chimaera expression in Shank3ΔC hippocampal neurons. Scale bar 10 μm. **C:** BRET variations between mGlu5-NLuc and Venus-Homer-GluN2B chimaera in Shank3ΔC neurons. Data are mean ± SEM of BRET recorded from at least 4 independent cultures. One-way ANOVA multiple comparison statistical analysis with Tukey correction to compare the last 2 time points of each curve. * indicates p-value < 0.05. Right: Long-lasting disruption calculated on the averaged last 2 indigo time points, expressed as a % of respective cyan time points. **D, E:** BRET variations in Shank3ΔC neurons expressing YEN (**D**) or AIMTOR (**E**) biosensors before (0’) and 12 min (12’) after depolarization. Scale bar 10 μm. Data are mean ± SEM of BRET intensities recorded in 8 to 30 neurons from 3 independent experiments. Point-by-point statistical tests and post hoc comparisons for **C, D and E** are indicated in Table 1.

### Repairing scaffolding complex dynamics rescues NMDA/AMPA excitatory postsynaptic currents ratio and improves cognitive impairments in Shank3ΔC mice

We next investigated whether repairing scaffolding complex dynamics with Homer-GluN2B chimaera could rescue cognitive impairments in Shank3ΔC mice, *in vivo*. We bilaterally injected P7 Shank3ΔC mice into the lateral ventricles with AAV encoding Homer-GluN2B chimaera tagged with a green fluorescent protein or only eGFP as a control. We also injected WT littermates with AAV expressing eGFP (Figure 5A). AAV-injection triggered a high transgene expression mainly in the hippocampus. Other structures like cortex, striatum, thalamus and subcortical nuclei are expressing the transgene with few heterogeneities among animals (Suppl. Figure 8). Importantly, Homer-GluN2B chimaera rescued Shank3ΔC synaptic transmission deficit, as shown by measurement of NMDA/AMPA excitatory postsynaptic currents ratio (Figure 5B). Of note, NMDA/AMPA ratio was not affected by the injection of the control AAV (Figure 3A and 5B), neither in WT nor in Shank3ΔC mice (p-value comparing mice injected or not with AAV ctrl: 0.4 in WT and 0.5 in Shank3ΔC; Mann et Whitney test), showing that surgery and viral expression had no effect *per se* on synaptic transmission. We further assessed the consequences on behaviors by performing a broad series of test in which Shank3ΔC mice are known to present deficits(7,8,10), from locomotion, stereotyped behaviors and anxiety. We also test Shank3ΔC in a fear conditioning paradigm. Compared to WT littermates, Shank3ΔC mice travelled less distance and with a lower speed in the open field test (Figure 5C). This aberrant locomotion was confirmed by their poor performance in the accelerating rotarod (Figure 5D) and not rescued by Homer-GluN2B chimaera expression. Shank3ΔC mice also present increased anxiety-like behaviors with a decreased time spent in the center during the open field test (Figure 5C), an increased time spent self-grooming (Figure 5E) and a higher score during fear conditioning (Figure 5F). The self-grooming test is also indicative of repetitive stereotyped behaviors in Shank3ΔC mice. Importantly, Homer-GluN2B chimaera expression in Shank3ΔC mice improved these anxiety-like and stereotyped behaviors in these three behavioral tests (Figure 5C, E and F). In the fear-conditioning test, Shank3ΔC mice tend to have a higher score compared to WT and Shank3ΔC Homer-GluN2B chimaera. This is the combination of a slightly lower freezing during habituation in day 2 ie contextual fear generalization and a higher amount of freezing during auditory cue presentation ie auditory CS+ fear response. In accordance with other studies, Shank3ΔC mice show no deficit in aversive learning, but display altered level of fear expression during the different phases of the test. This could be the result of a combination of an altered locomotion and an increased level of anxiety. While the marble-burying is usually used to test anxiety, it was recently described to test avoidance behavior to novelty in Shank3ΔC mice(7) who show no interest in burying marbles compared to WT littermates (Figure 5G). This behavior is unchanged in Shank3ΔC mice injected with Homer-GluN2B chimaera. Altogether, these experiments show that Shank3ΔC mice present deficits in locomotion, an avoidance behavior to novelty, anxiety-like behaviors and repetitive stereotyped behaviors. Injecting Homer-GluN2B chimaera in the lateral ventricles of Shank3ΔC mice decreased anxiety-like and repetitive stereotyped behaviors, meaning that a deficit in glutamate receptosome dynamics could be at the origin of the main core symptoms in ASD.

**Figure 5:**
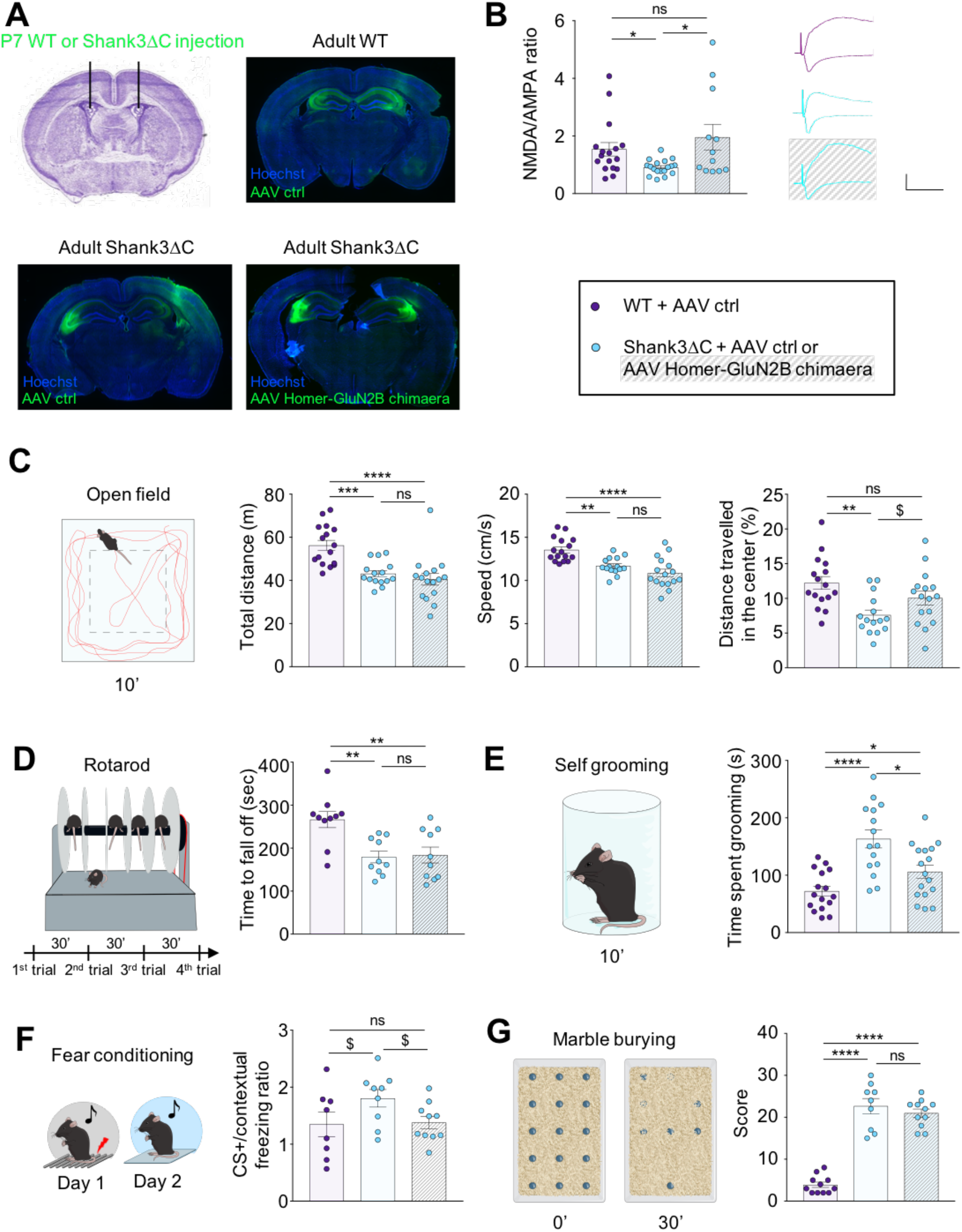
*in vivo* glutamate receptosome dynamics reestablishment rescues NMDA/AMPA currents and autistic-like behaviors in Shank3ΔC mice. **A:** Schema of injection and representative images of a coronal brain slice for each group. **B-G**: Data are mean ± SEM. ns: not significant; q-value: $ < 0.1; * < 0.05; ** < 0.01; *** < 0.001; **** < 0.0001; Kruskal-Wallis test with FDR post-test. **B:** Green CA1 pyramidal neurons from injected WT or Shank3ΔC mice were clamped at −80 and +40 mV to record AMPA and NMDA currents, respectively. n = 12-18 neurons from 5-7 mice per condition. Right panel: representatives average traces at −80 and +40 mV. Scale bar from the top to the bottom: 70, 60 and 40 pA over 50 ms. **C-G:** n = 8-17 mice per group. **C:** Open Field. Left panel: Top view of the arena with an illustration of a mouse trajectory during 10 minutes’ exploration. Dashed line square represents the center zone. Total distance and speed are measured during 10 minutes. Right panel: % of distance travelled in the center zone with respect to the total distance travelled during 10 minutes. **D:** Left panel: accelerating rotarod apparatus and protocol. Right panel: time in seconds to fall from the apparatus (average of the three last trials). **E:** Left panel: illustration of the self-grooming Plexiglas cylinder in which grooming was quantified. Right panel: time spent grooming by each group. **F:** Fear conditioning. Left panel: illustration showing the protocol. Right panel: Ratio of time spent freezing after CS+ over contextual time spent freezing. **G:** Marble Burying. Left panel: illustration showing the initial marble disposition and an example of buried marbles after 30 minutes. Right panel: Marble Burying Score quantification per group.

## Discussion

Shankopathies are one of the most common genetic causes of ASD(4,5). A deficit in NMDA neurotransmission represents a patho-physiological hallmark underlying cognitive impairments in Shankopathies and other ASD such as FXS. However, very little is known about the molecular synaptic mechanisms leading to this NMDA receptors impairment. In the past decade, we and others have highlighted the key role of mGlu5-Homer interaction in the control of synaptic plasticity and cognition. Indeed, following a first induction of NMDA-dependent plasticity, a decrease of mGlu5-Homer interaction is associated to physical interaction between mGlu5 and NMDA receptors and feed-back inhibition of NMDA currents, preventing subsequent induction of plasticity(21). In other words, scaffold remodeling controls synaptic availability for plasticity induction. Here, we used mGlu5-Homer interaction to report this synaptic availability by BRET. We depicted an activity-driven dynamic remodeling of mGlu5-Homer interactions, in WT mice. Transient neuronal depolarization, activating NMDA receptors to induce plasticity, causes a rapid break in the mGlu5-Homer interaction (within tens of seconds) while fundamental signals for synaptic plasticity expression, such as mTOR and ERK pathways are increased sustainably. In spines, this disruption is not reverted, possibly for hours. During this phase of plasticity expression, the scaffolding complex is broken and the NMDA receptors are blocked, making the synapse unavailable for subsequent plasticity induction. Indeed, a sustained increase in neuronal activity while mGlu5-Homer interaction is broken is no longer able to induce molecular and cellular plasticity. Interestingly, we observed the same kind of defects in the Shank3ΔC mouse model of ASD, in which a truncated form of Shank3 prevents scaffolding complex integrity. Shank3ΔC neurons indeed display a decreased basal NMDA/AMPA postsynaptic current ratio associated with a strong impairment in activity-driven increase of ERK and mTOR signaling pathways. We also confirmed previous reports, showing that these Shank3ΔC mice display strong anxiety-related and repetitive stereotyped behaviors(7,8,10). We then repaired the molecular synaptic complex in Shank3ΔC mice, while preserving its sensitivity to neuronal activity. Interestingly, this gene therapy restored the cellular plasticity and improved the cognitive performances of Shank3ΔC mice. Altogether, our data support a new theory according to which the glutamate receptosome complex, by its capacity to be reshaped by the neuronal activity, modifies NMDA receptors synaptic function and hence plays a decisive role in metaplasticity by sliding the induction threshold for plasticity. We further propose that molecular synaptic deficiencies would trigger synaptic plasticity impairments and atypical cognitive behaviors, reminiscent of ASD. This work opens opportunities of gene therapy strategies.

The first part of our study unveils mGlu5-Homer interaction dynamics in living cells upon neuronal activation. Neuronal depolarization induces a fast disruption of this interaction, uniformly in all neuronal sub-cellular compartments. However, the ability of the complex to re-form differs between sub-cellular compartments. In the soma, mGlu5 and Homer interact again as soon as the membrane potential resets back to resting potential. Successive transient depolarizations break and re-form the interaction, fleetingly. To date we know neither the molecular mechanisms sensing depolarization, nor the functional consequences of this ephemeral remodeling. Future studies will be required to better understand this phenomenon. By opposition, in half of the spines, the transient depolarization brakes mGlu5-Homer interaction within tens of seconds, in a long-lasting manner. We also show that NMDA receptors activation by neuronal depolarization is necessary to induce long-lasting disruption. The fast rupture between Homer and mGlu5 could be attributed, at least in part, to Ca^2+^ entry through NMDA receptors. Ca^2+^ would activate CamKIIα which can phosphorylate Homer long isoform, reducing its interaction with mGlu5(36). The immediate early gene Homer1a, which expression is induced by the increased neuronal activity, is then favored in the competition with the long form of Homer to interact in its place with the mGlu5 receptor(21). This could explain the long-lasting disruption of mGlu5-Homer interaction. This stably disrupted conformation of the receptosome could be a homeostatic mechanism to make synapses undergoing plasticity unavailable for further stimulation to induce new plasticity. We indeed show a strong heterogeneity in mGlu5-Homer dynamics in the spines that are attributable to differences in the basal interaction between these 2 proteins. The mGlu5-Homer disruption is almost absent in spines where the interaction is already weak. We can easily speculate that the BRET signal is low in these spines either because they are immature or because they recently underwent plasticity. Hence, the synaptic availability defined as the readiness of the synapse to a stimulus would be temporally blocked. This would favor the long-term expression of the initiated plasticity. These molecular dynamics at synapses explain reported roles of mGlu5-Homer in metaplasticity(37–39). Importantly, all these studies report Homer1a as the dominant-negative IEG breaking mGlu5-Homer interaction to allow synaptic plasticity. However, our experiments here demonstrate that mGlu5-Homer interaction is disrupted in few seconds while Homer1a synthesis takes about 20 minutes(40). This suggests that mGlu5-Homer interaction disruption could be the synaptic tag to attract Homer1a in potentiated synapses. This could be a homeostatic mechanism to lock a spine undergoing plasticity into a functional network, by decreasing temporarily its availability for plasticity induction. This hypothesis calls for further studies, to know whether and by which mechanisms a spine can be “unlocked” from a functional network.

The molecular cascade triggered by NMDA-dependent neuronal activation in physiological conditions includes scaffold disruption, increase of mGlu5 mobility, enhancement of mGlu5-NMDA clustering, inhibition of NMDA receptor activity and decrease of NMDA/AMPA ratio. These molecular events enable induction of ERK and mTOR activation. ERK activation is common to many forms of synaptic plasticity, in particular NMDA-dependent LTP in many brain areas(41,42). mTOR also sustains plasticity, memory storage, and cognition (see reviews(43,44)). While the important role of ERK and mTOR signaling pathways in neuronal plasticity has long been known, very little is known, however, concerning the activation kinetics of these signaling pathways due to the lack of real-time experimental approaches. Here we used mTOR and ERK biosensors(26,27) and showed that both kinases are immediately activated by transient increase of neuronal activity to reach a maximum 5 to 7 minutes later. Surprisingly, both ERK and mTOR substrates phosphorylation lasted for more than 20 minutes. These biosensors integrate kinases and phosphatases activities. This long-lasting substrate phosphorylation could report either kinases long-lasting activation or phosphatases inhibition during the late phase of neuronal plasticity. The latest hypothesis is supported by previous findings, showing that activation of mTOR is required only during the early phase of LTP(45). Interestingly, scaffold disruption prevents this activity-induced ERK and mTOR activation. This observation is coherent with our findings showing that scaffold disruption impairs activation of NMDA receptors(21) (Figure 2A) which are necessary to induce LTP via mTOR(45).

As discussed before, our data show sub-cellular differences in the capacity of mGlu5 and Homer to interact (soma) or not (spines) following a transient neuronal activation in wild type mice. Two non-exclusive hypotheses could explain this sub-cellular difference. First, Shank3 expression is confined to dendritic spines. Hence, counterintuitively, when mGlu5-Homer interaction is broken, Shank3 would be required to keep them apart from one another. Indeed, in the Shank3ΔC mouse model of ASD displaying a truncated form of Shank3, which cannot interact with Homer, mGlu5-Homer interaction dynamics in spines are reminiscent to the dynamics in the soma, i.e. a labile interaction highly sensitive to membrane potential and reversible at will. Second, NMDA receptors activation is required for long-lasting disruption of mGlu5-Homer interaction and NMDA/AMPA ratio is decreased in Shank3ΔC mice. Thus, a decrease of NMDA currents in Shank3ΔC mice could also account for impaired long-lasting disruption of Homer-mGlu5 interaction in spines. These 2 synergic mechanisms would explain the loss of long-lasting mGlu5-Homer disruption in Shank3ΔC mice. In favor of this assumption, we show that scaffold disruption in WT mice (using competitive TAT peptides) is sufficient to reduce by half the NMDA/AMPA ratio in CA1 pyramidal cells, corroborating a previous study showing a significant reduction of NMDA currents in similar conditions(34). Conversely, in Shank3ΔC mice, which display a higher basal endogenous mGlu5-NMDA interaction and lower NMDA/AMPA ratio, repair of the scaffolding complex and its sensitivity to neuronal activity using a chimeric protein is sufficient to restore the NMDA/AMPA ratio.

The chimeric protein also rescues a typical synaptic plasticity in Shank3ΔC mice. The present study demonstrates that mGlu5 receptosome integrity is required for induction of synaptic plasticity, and suggests the importance of the long-lasting disruption between mGlu5 and Homer for plasticity expression. Arguing in favor of this hypothesis, we previously showed that Homer1a expression was necessary to induce mGlu5-NMDA interaction and blockade of NMDA receptor activity after LTP induction (21). NMDA blockade prevents further change of synaptic excitability. We can easily speculate that Homer1a interaction with mGlu5 would prevent its re-association with the multimeric Homer and promote long-lasting disruption of the complex. The long-lasting disruption of the complex following transient depolarization is a common feature of native and restored complexes, enabling plasticity. Further studies will be required to incriminate the long-lasting disruption of the complex in plasticity expression.

In Shank3ΔC ASD model, NMDA/AMPA currents ratio is decreased compared to WT. The short-lived disruption of mGlu5-Homer interaction induced by transient depolarization have lost all control over the activity of glutamatergic receptors. Consistently, we show that ERK and mTOR signaling are not sensitive to the neuronal activity anymore. Interestingly, we observed an increase of the basal level of activated mTOR in Shank3ΔC neurons compared to WT neurons. A saturating activation of mTOR could explain the failure to increase this signaling pathway in Shank3ΔC neurons. Overactivation of mTOR has already been reported in other ASD models(27,46,47). Importantly, expression of the Homer-GluN2B chimaera to restore mGlu5 receptosome integrity in Shank3ΔC neurons was sufficient to set back mTOR basal level and allow its activation following transient depolarization. Our results show that glutamate receptosome remodeling is necessary to induce NMDA-dependent plasticity and activation of ERK and mTOR signaling pathways.

Autistic patients present two main symptoms: 1) deficits in social communication and 2) stereotyped behaviors meaning restricted and repetitive behavior, interest or activities. There are also numerous comorbidities such as anxiety, intellectual disability, cognitive deficits and avoidance behavior to novelty or motor control difficulties. Shank3ΔC mice phenotype recapitulates features of autistic patients. Among them, and as previously described, we found elevated anxiety, repetitive behaviors and decreased locomotor performances(7,8,10). The molecular rescue of glutamate receptosome dynamics using Homer-GluN2B chimaera efficiently rescued anxiety-like and repetitive stereotyped behaviors. Expression of anxiety has been associated with alteration of synaptic plasticity in the limbic circuit(48,49). Interestingly, the viral infection in both lateral ventricles early during development, has allowed a broad expression of the Homer-GluN2B chimaera in both dorsal and ventral hippocampi but also in other subcortical structures that are part of the limbic system such as striatum and amygdala. Based on our previous results, deficit in synaptic plasticity caused by the loss of glutamate receptosome dynamics would trigger anxiety related behaviors expression in Shank3ΔC. By opposition, Homer-GluN2B chimaera injection in the lateral ventricles of Shank3ΔC mice did not rescue locomotion nor avoidance behavior to novelty. This discrepancy presumably relies on the nature of brain areas targeted by the injection, and their involvement in specific behaviors. For example, knowing cerebellar contribution to locomotor behavior(50) and its relationship with approach/avoidance traits(51), the absence of transgene expression in the cerebellum (Suppl. Figure 8) is coherent with a lack of recovery of locomotion and avoidance behavior to novelty). Hence, this therapeutic gene therapy in mice opens promising opportunities to improve cognitive performances of ASD patients by targeting the glutamate receptosome dynamics. Besides, in agreement with a recent study performed in adults(52), this work supports promising opportunities to treat neurodevelopmental disorders by rescuing synaptopathies after birth.

To conclude, in this study, we showed that the reestablishment of glutamate receptosome integrity and dynamics in Shank3ΔC mouse model of ASD, using a chimaera, rescues normal functioning of glutamate receptors and activity-induced neuronal signaling. Importantly, this therapeutic strategy also relieves autistic behavioral features of mice, showing a causative link between glutamate receptosome alterations and cognitive behaviors in Shankopathies. Other Shankopathies are sensitive to the pharmacological regulation of mGlu5 receptor activity(53,54). Other ASD also relies on mGlu5-Homer interaction dysfunctions, such as fragile X syndrome(34,36) or phenylketonuria, a metabolic disorder that is characterized by autism and intellectual disability if untreated(55). Hence, the herein defined therapeutic strategy may be efficiently extended to these related pathologies.

## Acknowledgments

We thank the iExplore animal facility (IGF, Montpellier), CompAn behavioral phenotyping facility (MMDN, Montpellier) and the Arpege plateform (IGF, Montpellier) for the use of the Infinite F500 platereader for cell-population BRET. We thank Muriel Asari for illustrations. We thank Hélène Hirbec for helping with statistical analysis and Giorgio Cignitti for helping with patch on slices. We also gratefully acknowledge Paul F. Worley for the generous gift of the Shank3ΔC mouse model. This work was supported by the Agence Nationale de la Recherche (VC, ANR-18-CE16-0011-01) and the European Research Council (ERC) under the European Union’s Horizon 2020 research and innovation programme (JP, grant agreement No. 646788), ANR Lanthslider (JP, ANR-17-CE11-0046), Comitato Telethon Fondazione Onlus (grant no. GGP16131 to CV and GGP17176 to CS) and Regione Lombardia NeOn Progetto "NeOn" POR-FESR 2014-2020, ", ID 239047, CUP E47F17000000009 to CS and CV).

## Conflict of Interests

The authors declare no competing interests.

## Author contributions

E.M. and J.P. conceived research and wrote the manuscript; J.P., E.M., V.C., C.V., C.S, T.M., E.A., L.G., N.Bo. and S.S. designed research experiments; V.C., N.Bo. and F.G. cloned all plasmids. V.C. and E.M. produced viruses. E.M. performed cell population BRET experiments; E.M., J.P., V.S. and AL.H. performed microscopy BRET experiments. E.G. performed single particle training experiments. N.Be. analyzed single particle training experiments, with the help of E.G. and L.G. E.M. performed electrophysiology in slices, with the help of E.A. and F.R. V.C. performed co-immunoprecipitations with the help of E.M. and N.Bo. C.V., C.S. and F.G. designed and characterized the Homer-GluN2B construct. E.M. performed synaptosomes preparations. N.Bo. and E.M. performed immunohistochemistry on injected brains. S.S. performed surgery and viral injections. S.S. and J.A. performed behavioral experiments with the help of T.M. Y.C. developed toolsets for BRET analysis. J.P. supervised the project. All authors contribute to the preparation of the manuscript and approved it.

## Notes

### Competing Interest Statement

The authors have declared no competing interest.

