## Supplemental information for "Restoring Glutamate receptosome dynamics at synapses rescues Autism-like deficits in Shank3-deficient mice"

### Supplementary Methods

#### Synaptosomal preparations and WB analysis

Medium from DIV14-15 hippocampal cultures was replaced by KCl 3 mM. To observe the effects of a quick change of the neuronal resting potential on Homer and Homer1a expression, KCl 3 mM was replaced by KCl 55mM-evoked depolarization solution during 10 minutes. Then, medium was switched again for KCl 3 mM and cultures were put back in the incubator until 10, 20, 30, 60 or 120 minutes after the depolarization started. Then, cells were mechanically dissociated in ice-cold Syn-Per Synaptic Protein Extraction Reagent (Thermo scientific, reference 87793) supplemented with protease and phosphatase inhibitor cocktail (Thermo scientific), according to the manufacturer's protocol. Briefly, after a quick centrifugation to get rid of debris, all samples were then set at the same volume for the rest of the protocol. One equal volume was centrifuged for 20 minutes at 15000g. Pellets, that correspond to the synaptosomal part, were resuspended and the same volumes were loaded for each condition in the same gel. Proteins were eluted in Laemmli Sample buffer, resolved by SDS-PAGE on 4-15% gels (Bio-Rad), transferred onto nitrocellulose membranes and detected by immunoblot using the following primary antibodies: Homer1 rabbit (Synaptic systems, reference 160003, dilution 1:1000), GAPDH rabbit (Sigma-aldrich, G9545, dilution 1:25.000).

#### NMDA uncaging

NMDA uncaging was performed on DIV14 hippocampal neurons co-transduced at DIV6 with pAAV-CW3SL-mGlu5-NLuc and pAAV-CW3SL-Venus-Homer1C. BRET imaging was performed as described in the Single-cell BRET imaging section from Materials and methods excepted that a Plan Apochromat 40× /1.40 oil M27 objective was used. We sequentially exposed the donor 10 s and the acceptor 20 s and imaged 4 fields in each culture box; each field being recorded every 2 minutes. 1 mM photosensitive 4-methoxy-7-nitroindolinyl (MNI)-

caged-NMDA (Tocris) was applied concomitantly with Furimazine. After imaging of a 6 minutes baseline, NMDA was uncaged with violet LED (395/25 nm, 16 mW/mm$^2$), applying 4 pulses of 10 ms at 50 Hz on 2 over the 4 imaged fields of the culture box. Then, BRET was recorded for 12 minutes. As a second control, MNI-caged-NMDA was replaced by its vehicle (DMSO) and the same uncaging protocol was applied in all imaged fields of the culture box.

**Cell population BRET measurements**

Experiments were performed using the Infinite F500 (TECAN) 96-well plate-reader on DIV 14 to 16 neurons previously transduced at DIV6 with pWPT-Camk2αprom-mGlu5-NLuc and pWPT-Camk2αprom-Venus-Homer1C. BRET was measured in each well every 2 minutes (3 wells per condition in each experiment, pooled for the analysis). After a 10 minutes baseline, KCl 3 mM was replaced by KCl 3 mM alone or KCl 3 mM supplemented with TAT ctrl or TAT C-tail at a concentration of 50 μM.

**Single particle tracking**

We functionalized Qdot 655 ITK™ Amino PEG, (reference Q21521MP, Life Technologies) with a nanobody directed against mGlu5 as previously described(32). Briefly, 37 µl of QD655 were mixed with 13 µl of 30 mg/ml Bis[sulfosuccinimidyl] suberate (BS3, Sigma Aldrich) in 1M Borate buffer, pH 8. After 30 minutes of incubation and loading of this mixture onto a Nap-5 column (Life technologies), QD-BS3 conjugates were eluted with PBS pH 7.4. After a concentration step using a 50 kDa molecular weight cut-off filter (Life Technologies), QD-BS3 conjugates were incubated with 100 µl anti-mGlu5 neutral nanobodies (Fc-DN53, 1 mg/ml, a gift from Jean-Philippe Pin, IGF, Montpellier) for 2h with mild vortex. After 5-6 consecutives washing steps, the supernatant containing the QD-nanobodies conjugates was concentrated to 50 µl and unreacted BS3 sites were quenched by adding 2 µl of 1M Tris buffer pH 7.4.

DIV13-14 hippocampal primary neuronal cultures were incubated for 10 min with the QD-Nanododies complexes diluted at 1:1,000,000 in KCl 3 mM supplemented with 1% BSA to block non-specific binding and then washed. Neurons were transfected with DsRed-Homer as a synaptic marker. Detection, tracking and analysis of QDs for MSD and diffusion coefficients were performed as described previously(33). Multiple positions were imaged with an acquisition time of 50 ms with up to 500 consecutive frames for each position. Two baseline acquisitions at intervals of 5 minutes were performed before each treatment. The effect of a quick depolarization on mGlu5 surface diffusion was measured from 5 to 10 minutes after replacing KCl 3 mM with KCl 55 mM. Diffusion coefficients were calculated for every time-points of each microscopic field.

**HEK293 culture and transfection**

HEK293 cells were cultured in DMEM (Invitrogen) containing 10% FBS and antibiotics (Penicillin/Streptomycin) and transfected at 70–80% confluency (24 h after plating in 12-well plates with coverslips) using Lipofectamine 2000 (Invitrogen), or similar transfecting reagent, with the cDNA expression constructs (0.6 μg DNA per well). Transfected cells were fixed after for 36-48h with 4% paraformaldehyde supplemented with 4% sucrose and processed for the staining.

**Hippocampal primary cell culture transfection**

Rat hippocampal neuronal primary cultures were transfected using Lipofectamine 2000 at DIV11 with 2μg of cDNA for coverslips, then fixed with 4% paraformaldehyde supplemented with 4% sucrose and processed for the staining at DIV14–18. The specific shRNA for Shank3 was previously described and validated(1). The sequence is described in the Experimental

Procedure section RNA Interference and Relevant Plasmids. As scrambled control, we used the following sequence: GCTGAGCGAAGGAGAGAT.

**Immunocytochemistry**

Transfected rat hippocampal neuronal primary cultures and HEK293 cells were incubated with primary antibody in GDB [30 mM phosphate buffer, pH 7.4, 0.2% gelatin, 0.5% Triton X-100, 0.8 M NaCl (all Sigma–Aldrich)] for 3 h at room temperature. Cells were washed in 20 mM phosphate buffer containing 0.5 M NaCl and incubated with Cy3 and Cy5 conjugated secondary antibodies.

| **Fig 1C**; 2-way ANOVA, Tukey post-hoc | Baseline | | | | | First treatment | | | | | Second treatment | | | | | | | | | |
|---|---|---|---|---|---|---|---|---|---|---|---|---|---|---|---|---|---|---|---|---|
| Time points | 1 | 2 | 3 | 4 | 5 | 6 | 7 | 8 | 9 | 10 | 11 | 12 | 13 | 14 | 15 | 16 | 17 | 18 | 19 | 20 |
| KCl55+D-AP5-KCl3 — / KCl55-3 — | ns | ns | ns | ns | ns | ns | ns | ns | ns | ns | ** | *** | ** | *** | ** | ** | ** | ** | ** | ** |
| KCl3-3 — / KCl55+D-AP5-KCl3 — | ns | ns | ns | ns | ns | ** | ** | ** | *** | *** | * | ns | ns | ns | ns | ns | ns | ns | ns | ns |
| KCl55+D-AP5-KCl3 — / KCl55-55 — | ns | ns | ns | ns | ns | ns | * | * | * | ** | **** | **** | **** | **** | **** | **** | **** | **** | **** | **** |
| KCl3-3 — / KCl55-3 — | ns | ns | ns | ns | ns | **** | **** | **** | **** | **** | **** | **** | **** | **** | **** | **** | **** | **** | **** | **** |
| KCl55-3 — / KCl55-55 — | ns | ns | ns | ns | ns | ns | ns | ns | ns | ns | * | *** | **** | **** | **** | **** | **** | **** | **** | **** |
| KCl3-3 — / KCl55-55 — | ns | ns | ns | ns | ns | **** | **** | **** | **** | **** | **** | **** | **** | **** | **** | **** | **** | **** | **** | **** |

| **Fig 1D**; 2-way ANOVA, Tukey post-hoc | Baseline | | | | | First treatment | | | | | Second treatment | | | | | | | | | | Third treatment | | | | | Fourth treatment | | | | | | | | | | | | | | | | | | | |
|---|---|---|---|---|---|---|---|---|---|---|---|---|---|---|---|---|---|---|---|---|---|---|---|---|---|---|---|---|---|---|---|---|---|---|---|---|---|---|---|---|---|---|---|---|---|
| Time points | 1 | 2 | 3 | 4 | 5 | 6 | 7 | 8 | 9 | 10 | 11 | 12 | 13 | 14 | 15 | 16 | 17 | 18 | 19 | 20 | 21 | 22 | 23 | 24 | 25 | 26 | 27 | 28 | 29 | 30 | 31 | 32 | 33 | 34 | 35 | 36 | 37 | 38 | 39 | 40 | 41 | 42 | 43 | 44 | 45 |
| KCl3-3 — / KCl55-55 — | ns | ns | ns | ns | ns | ** | **** | **** | **** | **** | **** | **** | **** | **** | **** | **** | **** | **** | **** | **** | **** | **** | **** | **** | **** | **** | **** | **** | **** | **** | **** | **** | **** | **** | **** | **** | **** | **** | **** | **** | **** | **** | **** | **** | **** |
| KCl3-3 — / KCl55-3 — | ns | ns | ns | ns | ns | * | **** | **** | **** | **** | **** | **** | *** | **** | *** | ** | ** | *** | ** | ** | ** | ** | ** | *** | ** | ** | ** | ** | ** | ** | ** | ** | ** | ** | ** | *** | *** | ** | ** | ** | *** | ** | ** | * | *** |
| KCl3-3 — / KCl55-3-55-3 — | ns | ns | ns | ns | ns | * | **** | **** | **** | **** | **** | **** | **** | *** | **** | *** | ** | **** | ** | ** | **** | **** | **** | **** | **** | **** | **** | **** | **** | **** | *** | *** | **** | **** | **** | **** | **** | *** | **** | **** | ** | **** | *** | *** | *** |
| KCl55-3 — / KCl55-55 — | ns | ns | ns | ns | ns | ns | ns | ns | ns | ns | ns | ns | ns | * | ** | ** | ** | ** | ** | ** | ** | *** | ** | ** | *** | ** | ** | *** | ** | *** | *** | ** | ** | *** | ** | ** | ** | ** | ** | *** | * | ** | ** | ** | ** |
| KCl3-55 — / KCl55-3-55-3 — | ns | ns | ns | ns | ns | ns | ns | ns | ns | ns | ns | ns | ns | ** | * | * | ** | * | *** | *** | ns | ns | ns | ns | ns | ns | ns | ns | ns | * | ** | * | * | * | * | * | ** | ** | * | * | ** | * | ** | * | ** |
| KCl55-3 — / KCl55-3-55-3 — | ns | ns | ns | ns | ns | ns | ns | ns | ns | ns | ns | ns | ns | ns | ns | ns | ns | ns | ns | ns | ns | ** | ** | ** | *** | ns | ns | ns | ns | ns | ns | ns | ns | ns | ns | ns | ns | ns | ns | ns | ns | ns | ns | ns | ns |

| **Fig 1G**; one-way ANOVA, Dunnett post-hoc | KCl 3 mM | | | KCl 55 mM | | | | | KCl 3 mM | | | | | |
|---|---|---|---|---|---|---|---|---|---|---|---|---|---|---|
| Time points | 1 | 2 | 3 | 4 | 5 | 6 | 7 | 8 | 9 | 10 | 11 | 12 | 13 | 14 |
| Each time point is compared to time point 3 — | ns | ns | | **** | **** | **** | **** | **** | **** | **** | **** | *** | * | ns |

| **Fig 1H**; 2-way ANOVA, Sidak post-hoc | KCl 3 mM | | | KCl 55 mM | | | | | KCl 3 mM | | | | | |
|---|---|---|---|---|---|---|---|---|---|---|---|---|---|---|
| Time points | 1 | 2 | 3 | 4 | 5 | 6 | 7 | 8 | 9 | 10 | 11 | 12 | 13 | 14 |
| High basal BRET — / Low basal BRET — | *** | *** | *** | ns | ns | ns | ns | ns | ns | ns | **** | ns | ns | ** |

| **Fig 2E**; 2-way ANOVA, Sidak post-hoc | KCl 55 mM | | | KCl 3 mM | | | | | | | | | |
|---|---|---|---|---|---|---|---|---|---|---|---|---|---|
| Time points | 1 | 2 | 3 | 4 | 5 | 6 | 7 | 8 | 9 | 10 | 11 | 12 | 13 |
| TAT ctrl — / TAT C-tail — | ns | ns | ** | **** | **** | **** | **** | **** | **** | **** | **** | **** | **** |

| **Fig 2F**; 2-way ANOVA, Sidak post-hoc | KCl 55 mM | | | KCl 3 mM | | | | | | | | | |
|---|---|---|---|---|---|---|---|---|---|---|---|---|---|
| Time points | 1 | 2 | 3 | 4 | 5 | 6 | 7 | 8 | 9 | 10 | 11 | 12 | 13 |
| TAT ctrl — / TAT C-tail — | ns | ns | ns | *** | *** | *** | **** | **** | **** | **** | **** | *** | *** |

| **Fig 3B**; 2-way ANOVA, Sidak post-hoc | KCl 55 mM | | | KCl 3 mM | | | | | | | | | |
|---|---|---|---|---|---|---|---|---|---|---|---|---|---|
| Time points | 1 | 2 | 3 | 4 | 5 | 6 | 7 | 8 | 9 | 10 | 11 | 12 | 13 |
| WT — / Shank3ΔC ···· | ns | ns | ns | **** | ** | **** | **** | **** | **** | **** | *** | * | ** |

| **Fig 3C**; 2-way ANOVA, Sidak post-hoc | KCl 55 mM | | | KCl 3 mM | | | | | | | | | |
|---|---|---|---|---|---|---|---|---|---|---|---|---|---|
| Time points | 1 | 2 | 3 | 4 | 5 | 6 | 7 | 8 | 9 | 10 | 11 | 12 | 13 |
| WT — / Shank3ΔC ···· | ns | ns | * | **** | *** | *** | **** | * | ** | ** | * | ** | * |

| **Fig 3E**; 2-way ANOVA, Tukey post-hoc | Baseline | | | | | First treatment | | | | | Second treatment | | | | | | | | |
|---|---|---|---|---|---|---|---|---|---|---|---|---|---|---|---|---|---|---|---|
| Time points | 1 | 2 | 3 | 4 | 5 | 6 | 7 | 8 | 9 | 10 | 11 | 12 | 13 | 14 | 15 | 16 | 17 | 18 | 19 |
| WT KCl3-3 — / WT KCl55-55 — | ns | ns | ns | ns | ns | **** | **** | **** | **** | **** | **** | **** | **** | **** | **** | **** | **** | **** | **** |
| WT KCl3-3 — / WT KCl55-3 — | ns | ns | ns | ns | ns | **** | **** | **** | **** | **** | **** | **** | **** | **** | **** | **** | **** | **** | **** |
| WT KCl3-3 — / Shank3ΔC KCl3-3 ···· | ns | ns | ns | ns | ns | ns | ns | ns | ns | ns | ns | ns | ns | ns | ns | ns | ns | ns | ns |
| WT KCl3-3 — / Shank3ΔC KCl55-55 ···· | ns | ns | ns | ns | ns | **** | **** | **** | **** | **** | **** | **** | **** | **** | **** | **** | **** | **** | **** |
| WT KCl3-3 — / Shank3ΔC KCl55-3 ···· | ns | ns | ns | ns | ns | *** | **** | **** | **** | **** | **** | **** | ** | ** | ** | ** | ** | ns | * |
| WT KCl55-55 — / WT KCl55-3 — | ns | ns | ns | ns | ns | ns | ns | ns | ns | ns | ns | *** | **** | **** | **** | **** | **** | **** | **** |
| WT KCl55-55 — / Shank3ΔC KCl3-3 ···· | ns | ns | ns | ns | ns | *** | **** | **** | **** | **** | **** | **** | **** | **** | **** | **** | **** | **** | **** |
| WT KCl55-55 — / Shank3ΔC KCl55-55 ···· | ns | ns | ns | ns | ns | ns | ns | ns | ns | ns | ns | ns | ns | ns | ns | ns | ns | ns | ns |
| WT KCl55-55 — / Shank3ΔC KCl55-3 ···· | ns | ns | ns | ns | ns | ns | ns | ns | ns | ns | ns | * | *** | **** | **** | **** | **** | **** | **** |
| WT KCl55-3 — / Shank3ΔC KCl3-3 ···· | ns | ns | ns | ns | ns | *** | **** | **** | **** | **** | **** | **** | **** | **** | **** | *** | **** | **** | **** |
| WT KCl55-3 — / Shank3ΔC KCl55-55 ···· | ns | ns | ns | ns | ns | ns | ns | ns | ns | ns | ns | ns | ** | * | * | **** | ** | ns | **** |
| WT KCl55-3 — / Shank3ΔC KCl55-3 ···· | ns | ns | ns | ns | ns | ns | ns | ns | ns | ns | ns | ns | ns | ns | ns | ns | ns | ns | ns |
| Shank3ΔC KCl3-3 ···· / Shank3ΔC KCl55-55 ···· | ns | ns | ns | ns | ns | ** | **** | **** | **** | **** | **** | **** | **** | **** | **** | **** | **** | **** | **** |
| Shank3ΔC KCl3-3 ···· / Shank3ΔC KCl55-3 ···· | ns | ns | ns | ns | ns | ** | **** | **** | **** | **** | **** | ** | * | ** | * | ns | ** | * | ns |
| Shank3ΔC KCl55-55 ···· / Shank3ΔC KCl55-3 ···· | ns | ns | ns | ns | ns | ns | ns | ns | ns | ns | ns | ns | ** | ** | * | **** | ** | ** | **** |

| **Fig 3F**; 2-way ANOVA, Tukey post-hoc | Baseline | | | | | First treatment | | | | | Second treatment | | | | | | | | | | Third treatment | | | | | Fourth treatment | | | | | | | | | | |
|---|---|---|---|---|---|---|---|---|---|---|---|---|---|---|---|---|---|---|---|---|---|---|---|---|---|---|---|---|---|---|---|---|---|---|---|---|
| Time points | 1 | 2 | 3 | 4 | 5 | 6 | 7 | 8 | 9 | 10 | 11 | 12 | 13 | 14 | 15 | 16 | 17 | 18 | 19 | 20 | 21 | 22 | 23 | 24 | 25 | 26 | 27 | 28 | 29 | 30 | 31 | 32 | 33 | 34 | 35 | 36 |
| KCl3-3 ···· / KCl55-55 ···· | ns | ns | ns | ns | ns | **** | **** | **** | **** | **** | **** | **** | **** | **** | **** | **** | **** | **** | **** | **** | **** | **** | **** | **** | **** | **** | **** | **** | **** | **** | **** | **** | **** | **** | **** | **** |
| KCl3-3 ···· / KCl55-3 ···· | ns | ns | ns | ns | ns | *** | **** | **** | **** | **** | **** | *** | *** | * | ** | ** | *** | ns | ns | ** | * | ns | * | * | ns | ns | ns | ns | ns | ns | * | ns | ns | ns | ns | ns |
| KCl3-3 ···· / KCl55-3-55-3 ···· | ns | ns | ns | ns | ns | ** | **** | **** | **** | **** | **** | *** | **** | * | ** | * | *** | ns | * | *** | **** | **** | **** | **** | **** | **** | ** | *** | * | ** | * | ** | ns | ns | ns | * |
| KCl55-3 ···· / KCl55-55 ···· | ns | ns | ns | ns | ns | ns | ns | ns | ns | ns | ns | ns | ** | *** | * | ** | ** | *** | **** | **** | * | *** | ** | *** | **** | *** | **** | **** | *** | ** | ** | ** | * | *** | ** | **** |
| KCl55-55 ···· / KCl55-3-55-3 ···· | ns | ns | ns | ns | ns | ns | ns | ns | ns | ns | ns | ns | ns | ** | * | *** | ** | *** | ** | *** | ns | ns | ns | ns | ns | ns | ns | * | ** | * | ** | ns | * | **** | ** | ** |
| KCl55-3 ···· / KCl55-3-55-3 ···· | ns | ns | ns | ns | ns | ns | ns | ns | ns | ns | ns | ns | ns | ns | ns | ns | ns | ns | ns | ns | ns | ** | ** | ** | **** | * | * | ns | ns | ns | ns | ns | ns | ns | ns | ns |

| **Fig 3H**; one-way ANOVA, Dunnett post-hoc | KCl 3 mM | | | KCl 55 mM | | | | | KCl 3 mM | | | | | |
|---|---|---|---|---|---|---|---|---|---|---|---|---|---|---|
| Time points | 1 | 2 | 3 | 4 | 5 | 6 | 7 | 8 | 9 | 10 | 11 | 12 | 13 | 14 |
| Each time point is compared to time point 3 ···· | ns | ns | | *** | *** | **** | ** | ** | *** | ns | ns | ns | * | ns |

| **Fig 4C**; 2-way ANOVA, Tukey post-hoc | Baseline | | | | | First treatment | | | | | Second treatment | | | | | | | | |
|---|---|---|---|---|---|---|---|---|---|---|---|---|---|---|---|---|---|---|---|
| Time points | 1 | 2 | 3 | 4 | 5 | 6 | 7 | 8 | 9 | 10 | 11 | 12 | 13 | 14 | 15 | 16 | 17 | 18 | 19 |
| Shank3ΔC + chimaera KCl3-3 ···· / Shank3ΔC + chimaera KCl55-55 ···· | ns | ns | ns | ns | ns | *** | **** | **** | **** | **** | **** | **** | **** | **** | **** | **** | **** | **** | **** |
| Shank3ΔC + chimaera KCl3-3 ···· / Shank3ΔC + chimaera KCl55-3 ···· | ns | ns | ns | ns | ns | *** | **** | **** | **** | **** | **** | **** | **** | **** | ** | **** | ** | **** | **** |
| Shank3ΔC + chimaera KCl55-55 ···· / Shank3ΔC + chimaera KCl55-3 ···· | ns | ns | ns | ns | ns | ns | ns | ns | ns | ns | ns | ns | *** | **** | **** | **** | **** | *** | **** |

| **Fig 4D**; 2-way ANOVA, Tukey post-hoc | KCl 55/3 mM | | | KCl 3 mM | | | | | | | | | |
|---|---|---|---|---|---|---|---|---|---|---|---|---|---|
| Time points | 1 | 2 | 3 | 4 | 5 | 6 | 7 | 8 | 9 | 10 | 11 | 12 | 13 |
| Shank3ΔC KCl3-3 ···· / Shank3ΔC KCl55-3 ···· | ns | ns | ns | ** | ** | ** | ** | ** | ** | * | * | * | * |
| Shank3ΔC KCl3-3 ···· / Shank3ΔC + chimaera KCl3-3 | ns | ns | ns | ns | ns | ns | ns | ns | ns | ns | ns | ns | ns |
| Shank3ΔC KCl3-3 ···· / Shank3ΔC + chimaera KCl55-3 | * | *** | **** | **** | **** | **** | **** | **** | **** | **** | **** | **** | **** |
| Shank3ΔC KCl55-3 ···· / Shank3ΔC + chimaera KCl3-3 | ns | ns | ns | ** | ** | ** | ** | ** | *** | **** | ** | *** | ** |
| Shank3ΔC KCl55-3 ···· / Shank3ΔC + chimaera KCl55-3 | ns | ** | **** | **** | **** | **** | **** | **** | **** | **** | **** | **** | **** |
| Shank3ΔC + chimaera KCl3-3 / Shank3ΔC + chimaera KCl55-3 | ns | **** | **** | **** | **** | **** | **** | **** | **** | **** | **** | **** | **** |

| **Fig 4E**; 2-way ANOVA, Tukey post-hoc | KCl 55/3 mM | | KCl 3 mM | | | | | | |
|---|---|---|---|---|---|---|---|---|---|
| Time points | 1 | 2 | 3 | 4 | 5 | 6 | 7 | 8 | 9 |
| Shank3ΔC KCl3-3 ···· / Shank3ΔC KCl55-3 ···· | ns | ns | ns | ns | ns | ns | ns | ns | ns |
| Shank3ΔC KCl3-3 ···· / Shank3ΔC + chimaera KCl3-3 | **** | **** | **** | **** | **** | **** | **** | **** | **** |
| Shank3ΔC KCl3-3 ···· / Shank3ΔC + chimaera KCl55-3 | **** | ** | ns | ns | ns | ns | ns | ns | ns |
| Shank3ΔC KCl55-3 ···· / Shank3ΔC + chimaera KCl3-3 | **** | **** | **** | **** | **** | **** | **** | **** | **** |
| Shank3ΔC KCl55-3 ···· / Shank3ΔC + chimaera KCl55-3 | **** | **** | ns | ns | ns | ns | ns | ns | ns |
| Shank3ΔC + chimaera KCl3-3 / Shank3ΔC + chimaera KCl55-3 | ns | ns | ** | **** | **** | **** | **** | **** | **** |

| **Suppl. Fig 2A**; 2-way ANOVA, Tukey post-hoc | Baseline | | | | | Treatment | | | | | | | | | | | | | | |
|---|---|---|---|---|---|---|---|---|---|---|---|---|---|---|---|---|---|---|---|---|
| Time points | 1 | 2 | 3 | 4 | 5 | 6 | 7 | 8 | 9 | 10 | 11 | 12 | 13 | 14 | 15 | 16 | 17 | 18 | 19 | 20 |
| KCl3 + NMDA — / KCl55 + NMDA — | ns | ns | ns | ns | ns | ns | ns | ns | ns | ns | ns | ns | ns | ns | ns | ns | ns | ns | ns | ns |
| KCl3 + NMDA — / KCl3 — | ns | ns | ns | ns | ns | **** | **** | **** | **** | **** | **** | **** | **** | **** | **** | **** | **** | **** | **** | **** |
| KCl3 + NMDA — / KCl55 — | ns | ns | ns | ns | ns | ns | ns | ns | ns | ns | ns | ns | ns | ns | ns | ns | ns | ns | ns | ns |
| KCl55 + NMDA — / KCl3 — | ns | ns | ns | ns | ns | **** | **** | **** | **** | **** | **** | **** | **** | **** | **** | **** | **** | **** | **** | **** |
| KCl55 + NMDA — / KCl55 — | ns | ns | ns | ns | ns | ns | ns | ns | ns | ns | ns | ns | ns | ns | ns | ns | ns | ns | ns | ns |
| KCl3 — / KCl55 — | ns | ns | ns | ns | ns | **** | **** | **** | **** | **** | **** | **** | **** | **** | **** | **** | **** | **** | **** | **** |

| **Suppl. Fig 2B**; 2-way ANOVA, Tukey post-hoc | Baseline | | | Post-uncaging | | | | |
|---|---|---|---|---|---|---|---|---|
| Time points | 1 | 2 | 3 | 4 | 5 | 6 | 7 | 8 |
| Not uncaged MNI-NMDA — uncaged MNI-NMDA — | ns | ns | ns | ** | *** | **** | **** | **** |
| Not uncaged MNI-NMDA — "uncaged" vehicle — | ns | ns | ns | ns | ns | ns | ns | ns |
| Uncaged MNI-NMDA — "uncaged" vehicle — | ns | ns | ns | ns | *** | **** | **** | **** |

| **Suppl. Fig 4**; 2-way ANOVA, Tukey post-hoc | Baseline | | | | | Treatment | | | | | | | | | | | | | | | | | | | | | | | | | | | | |
|---|---|---|---|---|---|---|---|---|---|---|---|---|---|---|---|---|---|---|---|---|---|---|---|---|---|---|---|---|---|---|---|---|---|---|
| Time points | 1 | 2 | 3 | 4 | 5 | 6 | 7 | 8 | 9 | 10 | 11 | 12 | 13 | 14 | 15 | 16 | 17 | 18 | 19 | 20 | 21 | 22 | 23 | 24 | 25 | 26 | 27 | 28 | 29 | 30 | 31 | 32 | 33 | 34 |
| No TAT — / TAT ctrl — | ns | ns | ns | ns | ns | ns | ns | ns | ns | ns | ns | ns | ns | ns | ns | ns | ns | ns | ns | ns | ns | ns | ns | ns | ns | ns | ns | ns | ns | ns | ns | ns | ns | ns |
| No TAT — / TAT C-tail — | ns | ns | ns | ns | ns | ns | ns | ns | ** | *** | *** | ** | ** | *** | **** | *** | **** | ** | *** | *** | **** | **** | ** | *** | **** | *** | **** | **** | ** | **** | **** | **** | **** | *** |
| TAT ctrl — / TAT C-tail — | ns | ns | ns | ns | ns | ns | ns | ns | ns | ns | ns | ns | ns | ns | * | * | ns | ns | ns | ns | * | * | * | ** | ** | ** | *** | ns | ** | *** | ** | *** | *** | ** |

**Supplemental table 1: Statistical tests and post hoc comparisons.** Repetitive measures over time were compared using one-way ANOVA with Dunnett post-hoc when each time-point was compared to a control time-point, 2-way ANOVA with Sidak post-hoc when comparing the different time-points between two conditions and 2-way ANOVA with Tukey post-hoc when comparing every time-points of multiple conditions.

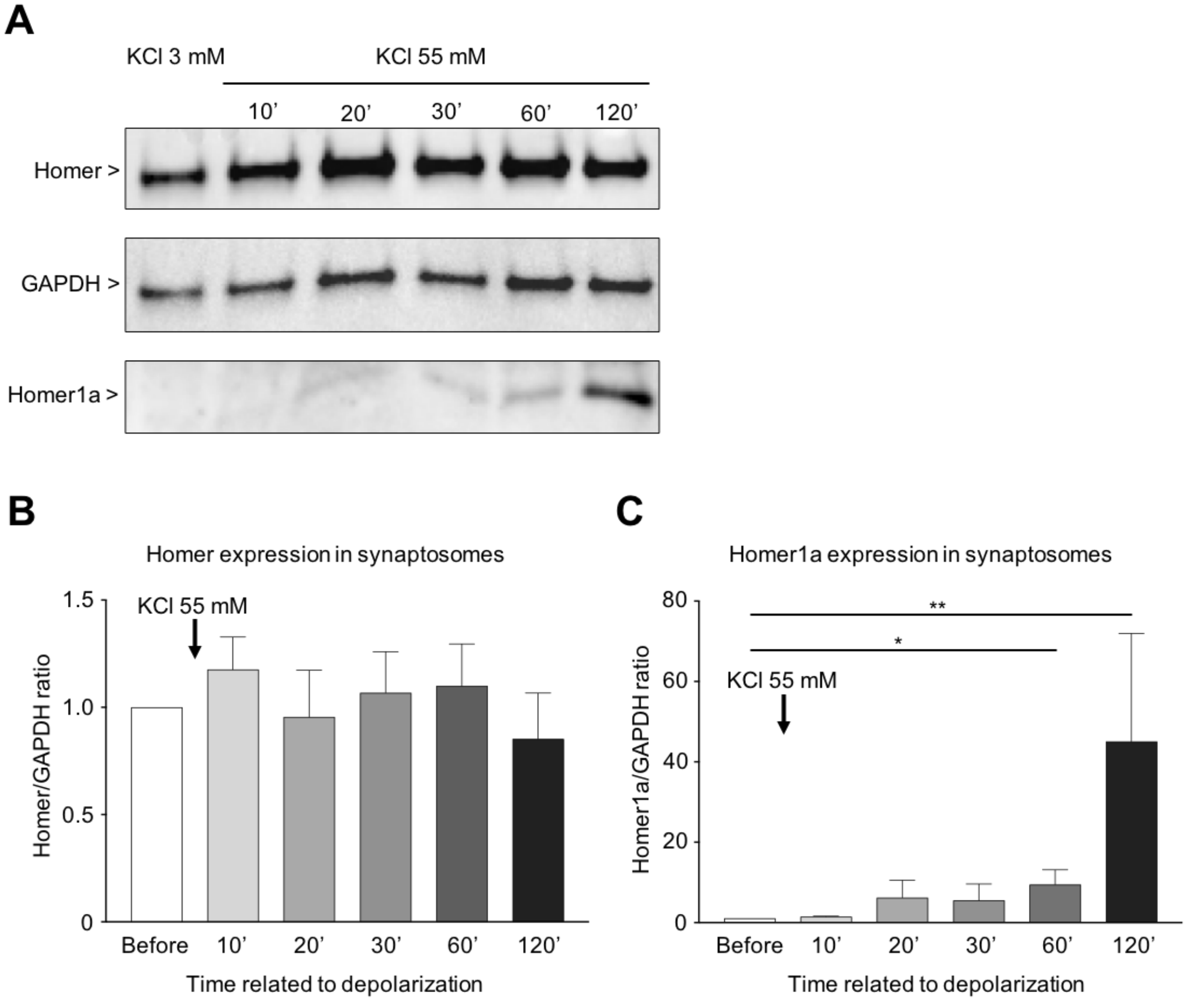


**Supplementary Figure 1: Homer1a expression significantly increases after transient neuronal activation. A:** Representative immunoblotting for Homer proteins on synaptosomal preparations from WT mature hippocampal cultures treated or not (KCl 3 mM) with KCl 55mM-evoked depolarization solution for 10 minutes and solubilized 10, 20, 30, 60 or 120 minutes after depolarization started. **B and C:** Histogram presents (**B**) Homer1 long form or (**C**) Homer1a short form expression, normalized by GAPDH expression, before and after different timings following a 10 minutes' depolarization. Data are mean ± SEM from 4 to 5 independent experiments. * indicates p-value $< 0.05$, ** $p < 0.01$; non-parametric Kruskal-Wallis test with Dunn's post-test.

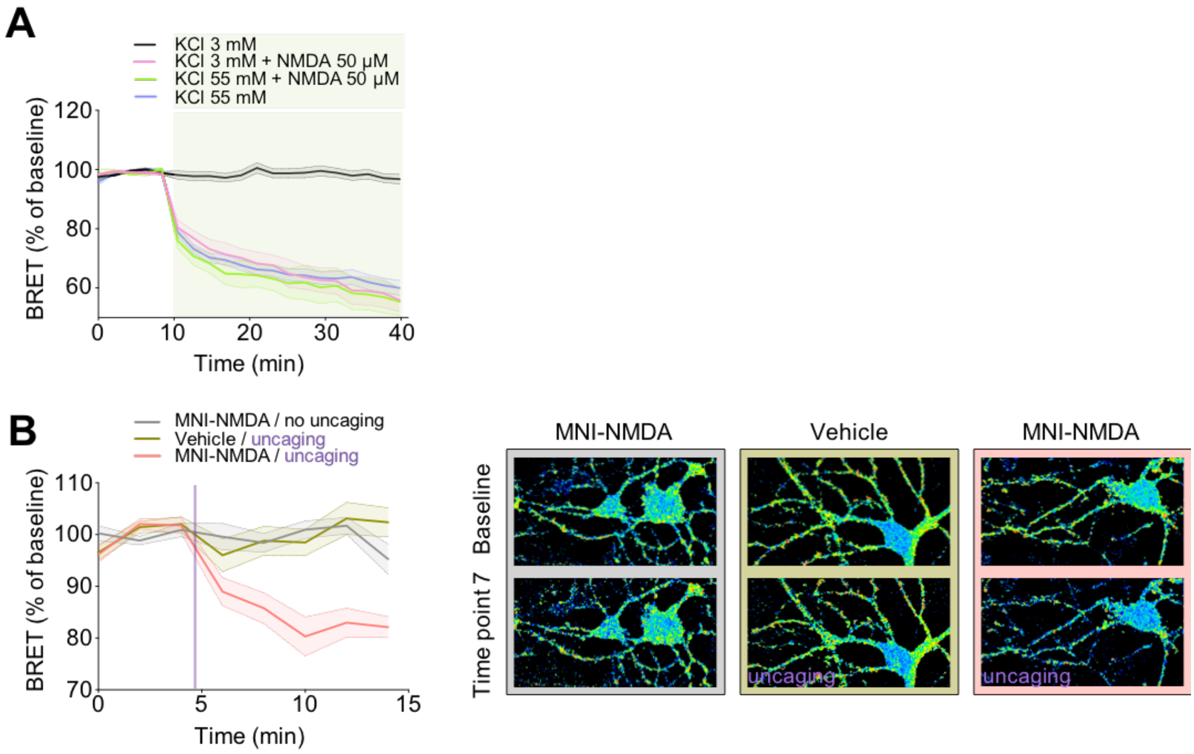


**Supplementary Figure 2: mGlu5-Homer interaction is disrupted by NMDA activation.** BRET recordings between mGlu5-NLuc and Venus-Homer in neurons from hippocampal cultures in cell population (**A**) or in microscopy (**B**). Real time measurements of BRET variations measured in neurons during changes of membrane potential with KCl 55 mM and/or during 50 µM NMDA application **(A)** or before/after NMDA uncaging **(B)**. Note that the effects of KCl 55 mM and NMDA are not additive (purple and green curves in **A**), suggesting that similar mechanisms are triggered by the two stimulations. Data are mean ± SEM of BRET intensities recorded in triplicate from at least 5 independent experiments (**A**) or on at least 15 fields from 2 independent experiments (we measured BRET from one to three neurons in each field) (**B**). Right **B** panel presents representative BRET images of the different conditions. Statistical tests and post hoc comparisons are indicated in Supplemental table 1.

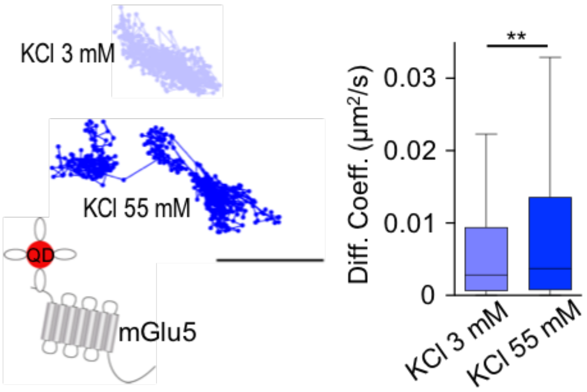


**Supplementary Figure 3: Transient depolarization induces an increase of mGlu5 mobility.**

mGlu5 single particle tracking in hippocampal neurons recorded in KCl 3mM and 55 mM (n = 12-15 neurons). **(**left) Representative trajectories (25 seconds), scale bar = 0.5 µm; (right) mGlu5 receptor total instantaneous diffusion coefficient ($\mu m^2 s^{-1}$) (n = 1171-1299 trajectories). Data are mean ± SEM; ** indicate p-value < 0.01; Mann-Whitney test.

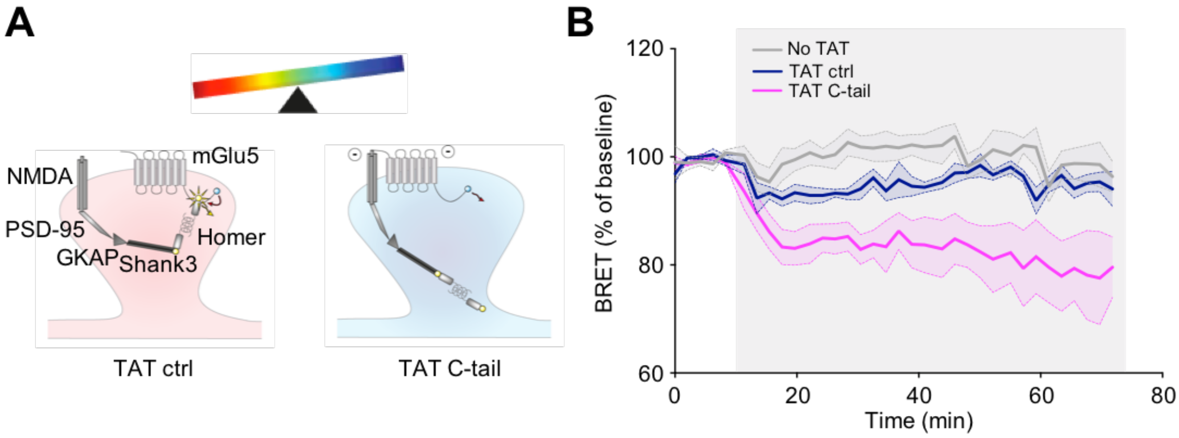


**Supplementary Figure 4: TAT C-tail disrupts mGlu5-Homer interaction. A:** Schematic illustration of the glutamate receptosome organization in spines depending on TAT C-tail or TAT ctrl presence. **B:** BRET variations between mGlu5-NLuc and Venus-Homer. Experiments were performed in KCl 3 mM. After a 10 minutes' baseline, medium was replaced with KCl 3 mM alone (grey) or KCl 3 mM supplemented with TAT ctrl (blue) or TAT C-tail (pink). Data are the mean ± SEM of BRET normalized to the baseline and recorded from 4 independent experiments. Statistical test and post hoc comparisons are indicated in the Supplemental table 1.

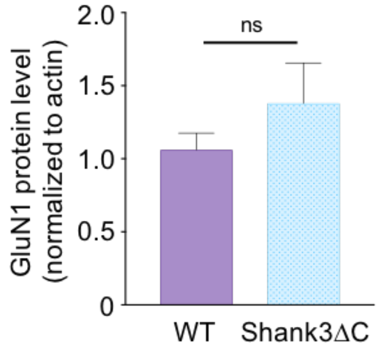


**Supplementary Figure 5: Total GluN1 protein level is stable in Shank3ΔC mice compared to WT littermates.**

Graph presents the GluN1 total lysate protein level normalized to actin expression from hippocampi of 7 Shank3ΔC mice and 7 WT littermates. See Co-immunoprecipitation section from Materials and methods for further details.

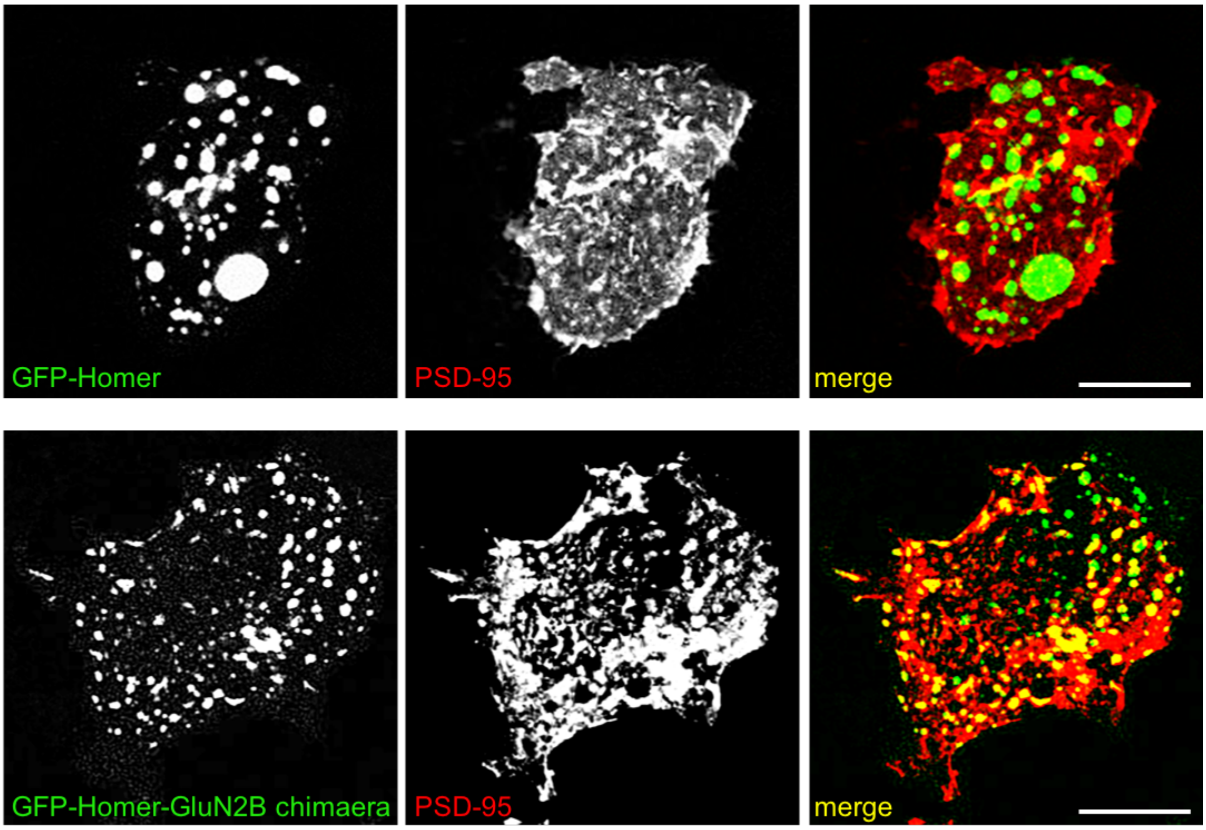


**Supplementary Figure 6: Homer-GluN2B chimaera co-localizes with PSD-95.** HEK293 cells were transfected with GFP-Homer or GFP-Homer-GluN2B plus PSD-95 as indicated in the panels. Only GFP-Homer-GluN2B forms intracellular co-clusters with PSD-95. Scale bar 5 µm.

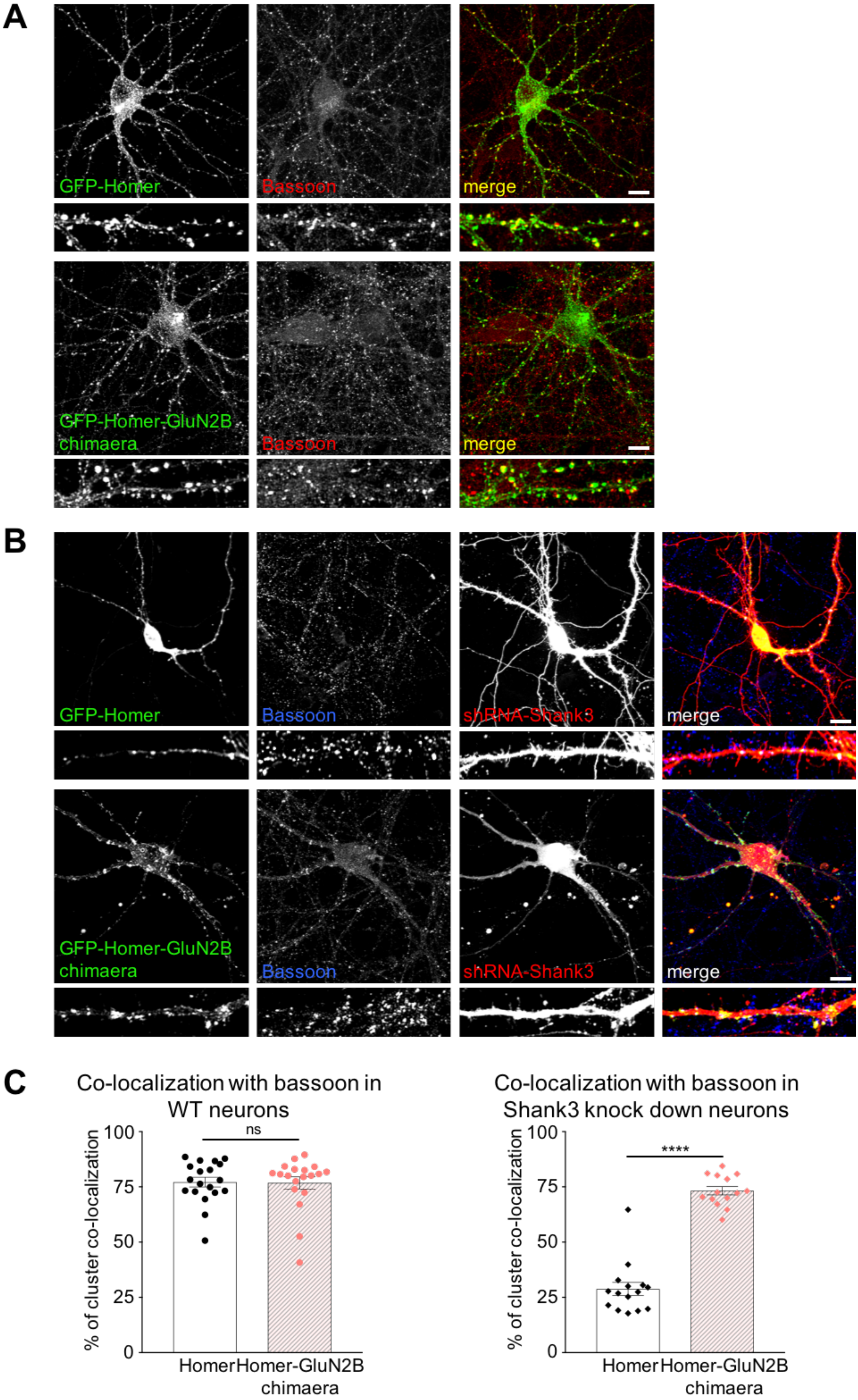
A
GFP-Homer
Bassoon
merge
GFP-Homer-GluN2B chimaera
Bassoon
merge
B
GFP-Homer
Bassoon
shRNA-Shank3
merge
GFP-Homer-GluN2B chimaera
Bassoon
shRNA-Shank3
merge
C
Co-localization with bassoon in WT neurons
% of cluster co-localization
100
75
50
25
0
ns
Homer
Homer-GluN2B chimaera
Co-localization with bassoon in Shank3 knock down neurons
% of cluster co-localization
100
75
50
25
0
****
Homer
Homer-GluN2B chimaera

**Supplementary Figure 7: Homer-GluN2B chimaera is correctly addressed in the absence of Shank3. A:** Rat hippocampal cultured neurons were transfected with GFP-Homer or GFP-Homer-GluN2B and immunolabeled with Bassoon antibody. **B:** Rat hippocampal cultured neurons were transfected with GFP-Homer or GFP-Homer-GluN2B plus a cDNA expressing shRNA for Shank3 and tdTomato an then immunolabeled with Bassoon antibody. **C:** The Quantification shows percent of colocalization between GFP-Homer or GFP-Homer-GluN2B clusters with Bassoon; 13 to 30 neurons were used for the quantification. Data are mean ± SEM from individual neurons; ns, **** indicate p-value > 0.05, < 0.0001, respectively; Mann-Whitney test. Scale bar 10 µm.

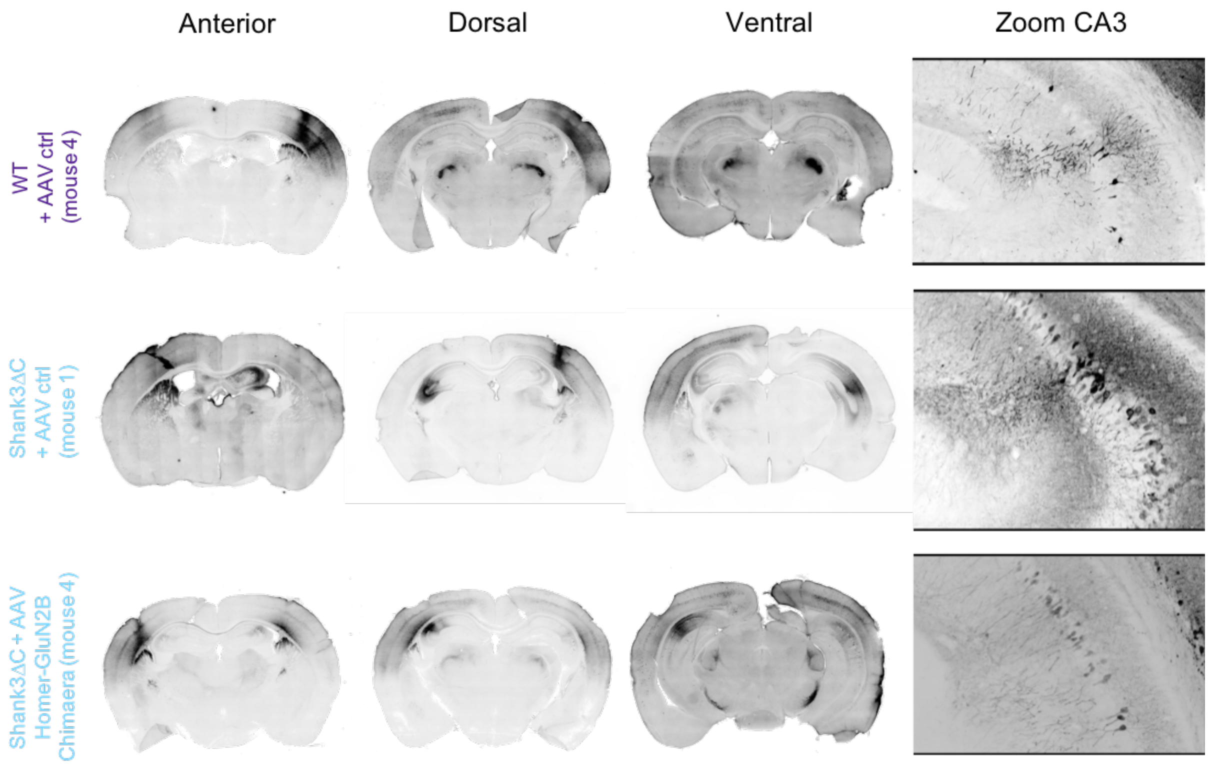


| | | Striatum | Cortex | | | Thalamus | Hippocampus | | | | | | Amygdala | Cerebellum |
|---|---|---|---|---|---|---|---|---|---|---|---|---|---|---|
| | | | | | | | CA1 | | CA3 | | DG | | | |
| | | | Ant | Mid | Post | | Dorsal | Ventral | Dorsal | Ventral | Dorsal | Ventral | | |
| WT + AAV ctrl | Mouse 1 | ++ | + | +++ | ++ | ++ | + | ++ | ++ | ++ | + | + | - | - |
| | Mouse 2 | + | - | + | ++ | + | ++ | +++ | ++ | ++ | + | + | - | - |
| | Mouse 3 | +++ | + | ++ | ++ | ++ | ++ | ++ | ++ | ++ | ++ | ++ | ++ | - |
| | **Mouse 4** | **+** | **-** | **++** | **+++** | **+** | **+** | **+** | **+** | **+** | **+** | **+** | **-** | **-** |
| | Mouse 5 | +++ | + | +++ | + | ++ | ++ | +++ | +++ | ++ | ++ | ++ | - | - |
| | Mouse 6 | + | - | +++ | ++ | + | ++ | +++ | +++ | + | + | + | + | - |
| | Mouse 7 | + | - | ++ | + | + | + | + | ++ | + | - | - | ++ | - |
| | Mouse 8 | - | - | + | + | ++ | +++ | +++ | +++ | +++ | +++ | +++ | - | - |
| | Mouse 9 | ++ | + | ++ | ++ | +++ | ++ | +++ | ++ | + | + | + | ++ | - |
| Shank3ΔC + AAV ctrl | **Mouse 1** | **++** | **+** | **++** | **+++** | **++** | **+** | **++** | **+++** | **++** | **+** | **++** | **++** | **-** |
| | Mouse 2 | ++ | ++ | +++ | +++ | +++ | + | +++ | ++ | ++ | ++ | + | + | - |
| | Mouse 3 | + | - | ++ | + | + | - | - | + | + | - | - | - | - |
| | Mouse 4 | +++ | +++ | +++ | +++ | ++ | + | + | ++ | +++ | + | + | +++ | - |
| | Mouse 5 | + | + | +++ | ++ | ++ | + | + | ++ | ++ | + | + | ? | - |
| | Mouse 6 | ++ | +++ | ++ | + | ++ | +++ | +++ | +++ | +++ | +++ | +++ | - | - |
| | Mouse 7 | +++ | ++ | +++ | +++ | +++ | ++ | +++ | +++ | ++ | ++ | ++ | - | - |
| | Mouse 8 | + | - | ++ | +++ | + | + | +++ | +++ | + | + | + | - | - |
| | Mouse 9 | +++ | + | +++ | +++ | ++ | +++ | +++ | +++ | +++ | ++ | ++ | ? | - |
| | Mouse 1 | - | - | + | + | - | ++ | ++ | +++ | +++ | - | - | - | - |
| | Mouse 2 | +++ | +++ | +++ | ++ | ++ | - | + | ++ | + | - | + | - | - |
| | Mouse 3 | ++ | ++ | +++ | ++ | +++ | +++ | +++ | +++ | +++ | +++ | +++ | + | - |
| | **Mouse 4** | **+++** | **+** | **+++** | **++** | **++** | **++** | **++** | **++** | **+** | **-** | **+** | **++** | **-** |
| | Mouse 5 | - | - | - | + | + | + | +++ | +++ | + | + | + | - | - |
| | Mouse 6 | +++ | +++ | ++ | +++ | +++ | - | - | - | - | - | - | + | - |
| | Mouse 7 | ++ | ++ | ++ | ++ | + | ++ | + | ++ | + | - | - | + | - |
| | Mouse 8 | ++ | ++ | ++ | ++ | - | - | + | +++ | + | - | - | - | - |
| | Mouse 9 | - | - | ++ | ++ | + | +++ | +++ | ++ | ++ | +++ | +++ | - | - |

**Supplementary Figure 8: Transgenes expression in WT and Shank3ΔC injected mice.** Representative illustrations of transgene expression in brain from mice who performed behavioral experiments. Slices chosen for representative images come from the mice in bold in the table. The table presents the intensity of expression by structure along the antero-posterior axis on a subsample of 9 mice per condition. Note that there are no signs of morphological disturbances of the cells (zoom).
